# Diurnal control of iron responsive element containing mRNAs through iron regulatory proteins IRP1 and IRP2 is mediated by feeding rhythms

**DOI:** 10.1101/2023.10.23.563593

**Authors:** Hima Priyanka Nadimpalli, Georgia Katsioudi, Enes Salih Arpa, Lies Chikhaoui, Alaaddin Bulak Arpat, Angelica Liechti, Gaël Palais, Claudia Tessmer, Ilse Hofmann, Bruno Galy, David Gatfield

**Author notes:** email addresses. equal contribution (Hima Priyanka Nadimpalli, Georgia Katsioudi, Enes Salih Arpa).

## Abstract

**Background:** Cellular iron homeostasis is regulated by iron regulatory proteins (IRP1 and IRP2) that sense iron levels (and other metabolic cues) and modulate mRNA translation or stability via interaction with iron regulatory elements (IREs). IRP2 is viewed as the primary regulator in liver, yet our previous datasets showing diurnal rhythms for certain IRE-containing mRNAs suggest a nuanced temporal control mechanism. The purpose of this study is to gain insights into the daily regulatory dynamics across IRE-bearing mRNAs, specific IRP involvement, and underlying systemic and cellular rhythmicity cues in mouse liver.

**Results:** We uncover high-amplitude diurnal oscillations in the regulation of key IRE containing transcripts in liver, compatible with maximal IRP activity at the onset of the dark phase. Although IRP2 protein levels also exhibit some diurnal variations and peak at the light-dark transition, ribosome profiling in IRP2-deficient mice reveals that maximal repression of target mRNAs at this time-point still occurs. We further find that diurnal regulation of IRE-containing mRNAs can continue in the absence of a functional circadian clock as long as feeding is rhythmic.

**Conclusions:** Our findings suggest temporally controlled redundancy in IRP activities, with IRP2 mediating regulation of IRE-containing transcripts in the light phase and redundancy, conceivably with IRP1, at dark onset. Moreover, we highlight the significance of feeding-associated signals in driving rhythmicity. Our work highlights the dynamic nature and regulatory complexity in a metabolic pathway that had previously been considered well-understood.

## Background

Mammalian physiology undergoes rhythmic changes orchestrated by the circadian system, which consists of hierarchically organised molecular oscillators found across most cell types (reviewed in [1]). The master clock in the brain’s suprachiasmatic nuclei entrains to the light-dark cycle and synchronizes peripheral oscillators through various cues that include hormonal signals, feeding-fasting cycles and body temperature rhythms. At the molecular level, the timekeeping mechanism is based on negative transcriptional feedback loops and drives daily cycles in gene expression (reviewed in [2]). Metabolism is a key rhythmically regulated process in organs such as the liver [3].

In addition to transcriptional regulation, various roles for post-transcriptional control of gene expression oscillations have emerged as well, involving in particular time-of-day-dependent changes in RNA stability and translation (reviewed in [4, 5]). Using ribosome profiling (ribo-seq) in mouse liver around-the-clock, mRNAs encoding ribosomal proteins have been identified as prominent examples of transcripts whose rhythmic regulation occurs specifically through translation [6, 7]. Moreover, in our previous work we reported that three transcripts encoding key proteins in iron metabolism, *Ferritin light chain 1 (Ftl1), Ferritin heavy chain 1 (Fth1)* and *Aminolevulinic acid synthase 2 (Alas2),* also show rhythmic translation [6].

More than 40 years ago, the discovery of the iron-responsive element, or IRE, occurred through studying how Ferritin mRNA (encoding a major iron storage protein) is translationally repressed and activated by low and high iron levels, respectively [8, 9]. *Alas2*, encoding the rate-limiting enzyme in heme biosynthesis in erythrocytes, was identified as IRE-containing a few years later [10]. Since then, several other IRE-containing transcripts have been identified and validated, and many more predicted ([11] and see below). These IREs are bound and regulated by IRPs (iron-regulatory proteins) in response to iron levels and other signals (O_2_, H_2_O_2_, NO) (reviewed in [12, 13]). The liver, a major iron storage organ, has been a key model for studying IRE/IRP regulation. Studies on mice lacking IRP1 (encoded by the *Aco1* gene) and IRP2 (encoded by *Ireb2*) have shown that IRP2 plays a major role in hepatic IRE regulation, with IRP1 having a limited role (e.g. [14–16]). Interestingly, none of these studies have explored a possible time-of-day component in this regulatory mechanism.

In this study, we systematically investigate the rhythmic regulation of IRE-containing transcripts beyond *Ftl1, Fth1,* and *Alas2*. We examine the dependence of this regulation on IRP1 and IRP2 using ribo-seq analysis on liver samples collected at defined timepoints during the light and dark phases from *Aco1/Irp1* and *Ireb2/Irp2* knockout mice. Additionally, we analyze existing datasets to determine whether rhythmic IRE regulation is primarily driven by the circadian clock or by feeding/fasting. Our findings prompt a reassessment of current models regarding IRP1 and IRP2 dependence of hepatic IRE regulation, a crucial mechanism for organismal iron balance.

## Results

### A subset of IRE-containing transcripts is rhythmically regulated in mouse liver

Our previous around-the-clock RNA-seq and ribo-seq datasets (Fig. 1A) had revealed hepatic rhythms in RNA abundance across >20 iron metabolic genes, as well as three transcripts – *Ftl1, Fth1* and *Alas2* – whose RNA levels were constant, yet they displayed robust rhythms in ribosome occupancy [6, 17]. *Ftl1, Fth1* and *Alas2* all contain an IRE in their 5’ untranslated region (5’ UTR) [11]. Iron regulatory proteins, IRP1 and IRP2, bind to such 5’ IREs to inhibit translation under conditions of low intracellular iron concentrations and as a function of other physiological cues; briefly, under high Fe(II), IRP1 assembles an iron-sulfur cluster and converts to ACO1 that is inactive for IRE binding, whereas IRP2 undergoes degradation (Fig. 1B) [11]. IRE sequences adapt characteristic hairpin structures (Supplementary Fig. S1A) that are crucial for IRP binding and have been used for IRE predictions [18, 19]. More than a dozen IRE-containing mRNAs have been reported in mammals, and we hence investigated whether rhythmic translation was more commonly found across other established 5’ IRE-containing transcripts. As compared to the robust translational rhythms on *Fth1, Ftl1* and *Alas2* in liver (Fig. 1C, upper), other mRNAs lacked similar high-amplitude oscillations in ribosome occupancy (Fig. 1D, upper). Thus, the translation of *Ferroportin* (*Slc40a1)* mRNA showed low-amplitude oscillations, and *Aconitase 2 (Aco2)* and *Hif2a/Endothelial PAS domain-containing protein 1 (Epas1)* were non-rhythmic, despite harbouring well-characterised 5’ IREs [20–22]. Note that although *Alas2* is considered to be a paralogue mainly expressed in erythrocytes, the observed rhythmic *Alas2* translation in liver is most likely of hepatocyte origin rather than a signal from red blood cells/reticulocytes within the liver sample, as liver single cell RNA-seq data [23] shows clear *Alas2* expression in hepatocytes, see Supplementary Fig. S2). We next compared the liver profiles with those from kidney, which revealed that *Ftl1, Fth1* and *Slc40a1* showed blunted ribosome occupancy rhythms in this organ (Fig. 1C-D, lower). *Aco2* and *Epas1* were clearly non-rhythmic in kidney. We considered it a plausible hypothesis that distinct sequence/structural features of the IREs could underlie the observed presence or lack of rhythmicity. However, despite differences across IREs with regard to the length of the paired stem, the positions of bulge nucleotides, or IRE location relative to mRNA 5’ end or start codon, no particular feature appeared to distinguish rhythmicity-associated from other IREs (Supplementary Fig. S1B-C).

**Figure 1.**
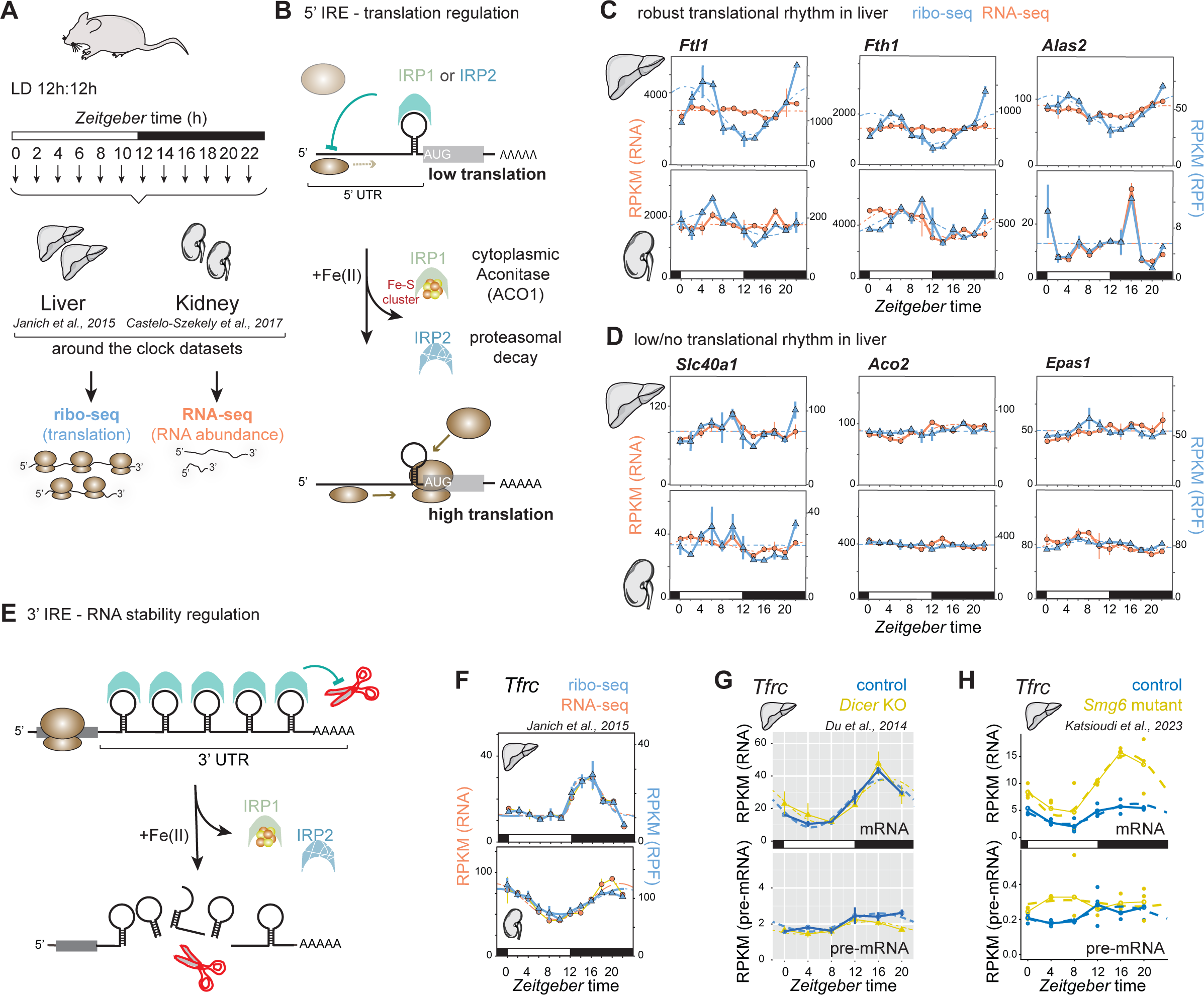
Transcript- and tissue-specific rhythmicity of IRE-containing mRNAs. (**A**) Schematic of the dataset collection carried out for the studies [6, 17] for the liver and kidney gene expression graphs shown in panels C and D (below). (**B**) Schematic model of IRP activity on 5’ IREs. IRP1 and IRP2 bind to IRE hairpins under conditions of low iron, thereby inhibiting scanning ribosomes from accessing the start codon. When iron is abundant, IRP1 assembles an Fe-S cluster, converting it to cytoplasmic Aconitase (ACO1) that is unable to bind IREs. IRP2 is degraded in an iron-dependent proteasomal pathway that relies on an iron-sensor F-box protein, FBXL5. After removal of IRPs from the 5’ IRE sequence, ribosomes can translate the coding sequence (CDS). (**C**) RPKM values of CDS-mapping ribosome protected fragment (RPF) data (blue) and RNA-seq data (orange) from the datasets of [6, 17] for the transcripts *Ftl1, Fth1* and *Alas2* around the 24-h daily cycle in liver (upper) and kidney (lower). All three genes show high amplitude translation in liver. Means per timepoint are plotted; “error bars”/vertical lines connect the two biological replicates. Dashed lines represent rhythmic curve fittings to the data, as described in [6]. (**D**) As in panel (C), but for the non-/less-rhythmically translated IRE-containing transcripts *Slc40a1/Ferroportin, Aco2, Epas1/Hif2-alpha*. (**E**) For 3’ UTR IREs, the binding of IRPs inhibits access to endonucleases (depicted as red scissors) that would otherwise initiate transcript degradation. When IRPs dissociate due to the mechanisms described in panel (B), the mRNA is destabilised and becomes less abundant. *Tfrc* mRNA is the best known example of this kind and contains five IREs. (**F**) RPKM values of CDS-mapping ribosome protected fragment (RPF) data (blue) and RNA-seq data (orange) from the datasets of [6, 17] for the *Tfrc* transcript around the 24-h daily cycle in liver (upper) and kidney (lower). Means per timepoint are plotted; “error bars”/vertical lines connect the two biological replicates. Dashed lines represent rhythmic curve fittings to the data, as described in [6]. (**G**) Expression plots (RPKM) showing around-the-clock data from [26] for *Tfrc* mRNA (exonic reads; upper panel) and pre-mRNA (intronic reads; lower panel) in control (blue) and miRNA-deficient (*Dicer* KO) livers (yellow). Means per timepoint are plotted; “error bars”/vertical lines connect the two biological replicates. The two time series are plotted together in the same graph (vertical lines connect the data points from the two series). Rhythms are seen for mRNA but not for pre-mRNA, indicating that oscillations are post-transcriptionally generated (RNA stability). Dashed lines represent rhythmic curve fittings to the data, as described in [26]. (**H**) Expression plots for around-the-clock RNA-seq data from [27] for *Tfrc* showing mRNA (top panel; exonic reads) and pre-mRNA (bottom panel; intronic reads) for *Smg6* mutants (yellow) and controls (blue). RPKM values of individual mouse livers are shown as dots with solid lines connecting the means across time points. Dashed lines represent rhythmic curve fittings to the data, as described in [27].

In some cases, IREs are located in 3’ UTRs. In *Transferrin receptor (Tfrc*) mRNA, multiple 3’ IREs regulate transcript stability in an IRP-dependent manner, involving endonucleolytic cleavage (Fig. 1E; Supplementary Fig. S1D) [24]. We hence analysed diurnal *Tfrc* expression profiles. *Tfrc* mRNA abundance was indeed robustly rhythmic in liver and kidney, and ribosome footprints closely matched the mRNA profiles, indicating that no additional rhythmicity was imparted at the level of translation (Fig. 1F). We then sought to evaluate if the observed *Tfrc* mRNA rhythms were of post-transcriptional origin, in line with the 3’ IRE mechanism and previous observations [6, 25]. To this end, we leveraged two around-the-clock mouse liver datasets [26, 27] in which we had carried out RNA-seq on total RNA, allowing the quantification of both unspliced pre-mRNA (intron-mapping reads) and of mature mRNA (exon-mapping reads). With pre-mRNA levels serving as a proxy for nascent RNA/transcription at the *Tfrc* locus, both datasets confirmed that *Tfrc* was transcribed in a non-rhythmic fashion, yet its mRNA accumulated rhythmically around-the-clock (Fig. 1G-H, data from control mice). We concluded that *Tfrc* rhythms were generated post-transcriptionally by modulating transcript stability in a diurnal fashion; this finding is further supported by a recent preprint [28].

As a side note, one of the analysed datasets (Fig. 1G) was part of a study that contained a matching cohort of animals with a disrupted miRNA biogenesis pathway (hepatocyte-specific deletion of *Dicer* [26]). It has previously been proposed that *Tfrc* mRNA stability is governed by the interplay between IRPs and miR-7/miR-141 binding at the 3’ IRE stem-loops [29]. Given that *Tfrc* mRNA profiles were unaltered in *Dicer* knockouts (Fig. 1G), we deem it unlikely that miRNA-mediated mechanisms make strong contributions to hepatic 3’ IRE regulation. The other analysed dataset (Fig. 1H) included an experimental cohort that carried a hepatocyte-specific mutation within the endonuclease that catalyse RNA degradation in the nonsense-mediated mRNA decay (NMD) pathway, *Smg6* [27]. Briefly, the increase in *Tfrc* mRNA in *Smg6^mut/mut^* livers (Fig. 1H) is most likely the result of a stabilization of several *Tfrc* mRNA variants that contain retained introns (see ENSEMBL annotations, locus ENSMUSG00000022797) and that are usually degraded through the NMD pathway.

### Rhythmic translation/mRNA abundance predicts phases of maximal and minimal IRP activity

From the above analyses we concluded that a subset of IRE-containing transcripts underwent pronounced rhythmic regulation in mouse liver. Of these, the 5’ IRE transcripts showed similar rhythmic phase, with a peak in translation around ZT4-6 and a trough at ∼ZT12 (Fig. 1C). Given that IRP-IRE interaction inhibits translation (Fig. 1B), we would hence expect IRP activity on these transcripts to be minimal at ZT4-6 and maximal at ZT12. For *Tfrc*, IRP binding is expected to increase mRNA stability and we would predict maximal IRP activity during the rising phase of *Tfrc* mRNA abundance, i.e. at ∼ZT12 as well. Therefore, the ensemble of observed rhythms would be consistent with a mechanism involving time-of-day-dependent IRP activity on IREs that is minimal and maximal at ZT4-6 and ZT12, respectively (Fig. 2A). Diurnal regulation could be mediated simply by daily changes in IRP protein abundance. We had dismissed this mechanism, however, in our previous work as we had observed no rhythmicity (or, at most, very modest daily changes) for liver IRP1/ACO1 or IRP2/IREB2 at the level of mRNA, ribosome footprints or protein [6]. Since publication of our study it has been reported that the antibody we had used to quantify IRP2/IREB2 (abcam ab1811539) not only detects IRP2/IREB2 (as marketed by the vendor at the time), but also IRP1/ACO1 [30]. Therefore, our original analysis of around-the-clock IRP2 protein levels was undoubtedly flawed, representing an overlay of both IRPs rather than IRP2 specifically. Moreover, it has been reported that in colon cancer cells, the *Ireb2* gene is circadianly transcribed through BMAL1:CLOCK, engendering *Ireb2* mRNA and IRP2/IREB2 protein oscillations that subsequently regulate *Tfrc* stability in a rhythmic fashion [31]. The ensemble of these findings prompted us to revisit the temporal dynamics of IRP expression across our datasets. Thus, our RNA-seq and ribo-seq data showed modest rhythmicity (<1.5-fold peak-to-trough amplitudes) for *Aco1* and *Ireb2* in liver but not in kidney (Fig. 2B). For *Ireb2*, the mRNA phase – maximal abundance around ZT10 – was in principle compatible with the BMAL1:CLOCK regulation reported from colon cancer cells [31]. However, the shallow peak-trough amplitudes that *Ireb2* displays for both RNA and pre-mRNA (Fig. 2C, lower) indicated that rhythmic transcriptional activation through BMAL1:CLOCK would not appear to be an important driver of *Ireb2* mRNA synthesis in liver. For comparison, in the same dataset *Arntl/Bmal1* itself, and well-established BMAL1:CLOCK targets such as *Dbp* [32], present very high amplitude mRNA and pre-mRNA oscillations (Fig. 2D).

**Figure 2.**
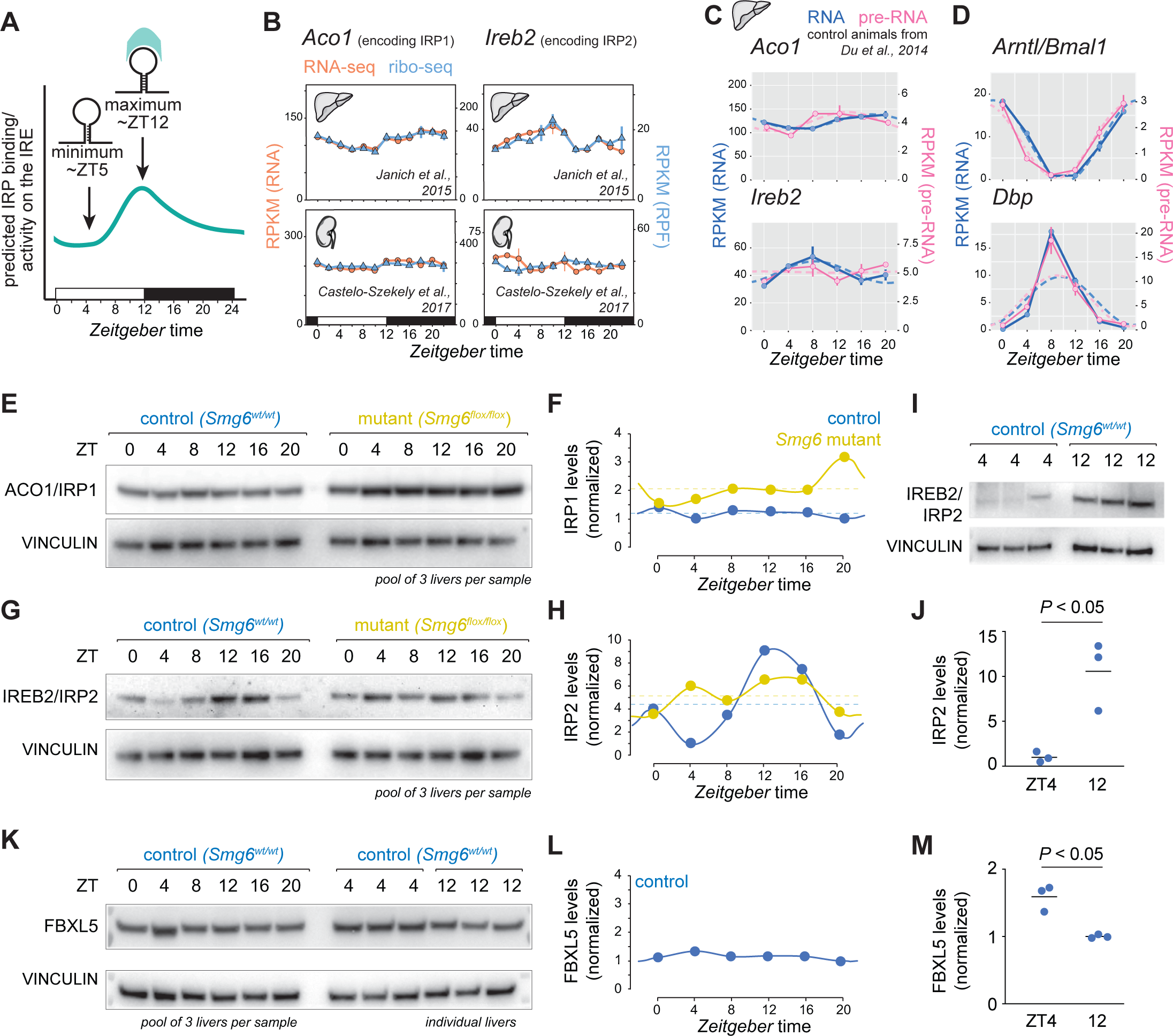
Rhythmic regulation is consistent with diurnal IRP activity and low and high IRP2 abundance at ZT5 and ZT12. (**A**) Schematic representation of the model deduced from the gene expression data from Fig. 1C (rhythmic 5’ IREs for *Ftl1, Fth1* and *Alas2*) and Fig. 1F-H (rhythmic 3’ IRE for *Tfrc*). The observed rhythms are consistent with minimal and maximal IRP activity at the light-phase (ZT5) and light-dark transition (ZT12) timepoints, respectively. (**B**) As in Fig. 1C, RPKM values of CDS-mapping ribosome protected fragment (RPF) data (blue) and RNA-seq data (orange) from the datasets of [6, 17] for the transcripts *Aco1* (encoding IRP1) and *Ireb2* (encoding IRP2) around the 24-h daily cycle in liver (upper) and kidney (lower). Means per timepoint are plotted; “error bars”/vertical lines connect the two biological replicates. Dashed lines represent rhythmic curve fittings to the data, as described in [6]. (**C**) Expression plots (RPKM) showing around-the-clock data from [26] for the indicated transcripts, *Aco1* and *Ireb2,* for mRNA (exonic reads; blue) and pre-mRNA (intronic reads; magenta) in control livers. The data from the miRNA-deficient (*Dicer* KO) livers were ommitted from these analyses. Means per timepoint are plotted; “error bars”/vertical lines connect the two biological replicates. Dashed lines represent rhythmic curve fittings to the data, as described in [26]. (**D**) As in (C), but for transcripts *Arntl/Bmal1* and *Dbp*. (**E**) Western blot analysis of total liver proteins (pool of three mice per sample), for ACO1/IRP1 and VINCULIN (loading control), from control mice (blue; on left side) and *Smg6* mutant mice (yellow; on right side). Same samples as in [27] were used. (**F**) Quantification of Western blot from (E). ACO1/IRP1 itensities were normalised to control band intensities (VINCULIN). (**G**) Western blot analysis of total liver proteins as in (E), for IREB2/IRP2 and VINCULIN (loading control). (**H**) Quantification of Western blot from (G). IREB2/IRP2 itensities were normalised to control band intensities (VINCULIN). (I) Western blot analysis of total liver proteins from individual animals for IREB2/IRP2; the same extracts that constituted the pools ZT4 and ZT12 (control series) in panel G were used. (J) Quantification of Western blot from (I). IREB2/IRP2 itensities were normalised to control band intensities (VINCULIN). Statistical significance evaluated via Student’s t-test. (K) Western blot analysis for FBXL5 and VINCULIN (loading control) in total liver proteins from the control series samples from [27]. Left side of blot: around-the-clock, pool of three mice per sample; right side: individiual mice, as in panel (I). (L) Quantification of around-the-clock data from Western blot from (K). FBXL5 itensities were normalised to control band intensities (VINCULIN). (M) Quantification of Western blot from individual livers at ZT4 and ZT12 from (K), right part of blot. FBXL5 itensities were normalised to control band intensities (VINCULIN). Statistical significance evaluated via Student’s t-test.

The main regulation of IRP1 activity and of IRP2 abundance occurs through post-translational mechanisms (Fig. 1B) [13]. Given the issues with antibody specificity described above, we re-evaluated IRP1 and IRP2 protein levels around-the-clock using highly specific monoclonal antibodies that are free of cross-reactivity (see later, Fig. 4G, J). Using a time series of total liver protein extracts from the same animals that showed *Tfrc* mRNA rhythms in Fig. 1H [27], Western blot analysis confirmed that ACO1/IRP1 showed constant abundance (Fig. 2E, F). By contrast, the Western blot for IREB2/IRP2 revealed high-amplitude rhythmic protein accumulation in the control liver time series, with minimal and maximal protein abundances around ZT4 and ZT12, respectively (Fig. 2G, H). The difference in IRP2 protein levels quantified from single liver samples at trough (ZT4) and peak (ZT12) showed an ∼10-fold amplitude with some inter-individual variability (Fig. 2I, J). Of note, we also included our previous study’s “experimental condition” samples (*Smg6^mut/mut^*[27]) in our Western blots. Interestingly, IRP2 protein rhythm was blunted in *Smg6^mut/mut^* livers (Fig. 2G, H), despite *Tfrc* mRNA oscillating with high amplitude according to RNA-seq analysis from exactly the same tissue samples (Fig. 1H) – a discrepancy that led us to conclude that there was no simple, one-to-one correspondence between oscillations in IRP2 protein abundance and IRE-containing transcript regulation. We also analysed the levels of FBXL5, which is the F-box protein that targets IREB2/IRP2 for proteasomal degradation and is key for the regulation of IRP2 stability and abundance. According to the established mechanism, FBXL5 acts as an iron sensor that is stabilised by Fe binding under high iron conditions, yet unfolded and degraded during iron scarcity [33, 34]. In addition, an FBXL5-independent mechanism of IREB2/IRP2-dependent target regulation has recently been identified as well [35]. Our Western blot analysis in the control liver time series and in single liver samples indicated that FBXL5 protein levels showed modest rhythms only (Fig. 2K-M). We therefore deemed it unlikely that the observed strong diurnal changes in IREB2/IRP2 abundance (Fig. 2G, H) would be the simple outcome of FBXL5 abundance changes (Fig. 2K-M). Taken together, our analyses revealed hepatic rhythms in IRP2 abundance that have previously gone unnoticed. Still, the lack of clear correspondence between rhythmic patterns seen for the abundances of IRP2 and FBXL5 proteins as well as *Tfrc* mRNA, highlight the regulatory complexity within the IRP/IRE system.

### In vivo recording of diurnal gene expression oscillations shows unperturbed circadian mPER2::LUC reporter rhythms in Aco1 and Ireb2 knockout mice

Two reasons pointed to a possible involvement of IREB2/IRP2 in diurnal IRE regulation. First, the observation that – in the time series data shown in Fig. 2G, H – IRP2 did show rhythmicity with lower and higher abundances in the light and dark phases, respectively. Second, IRP2 can be considered the main IRP activity in liver, given that changes to IRE regulation have generally been reported upon IRP2 deficiency rather than IRP1 loss [14, 36]. However, if IRP1 does not constitute a major IRP activity in liver, it is intriguing that the protein is particularly abundant in this organ, as also confirmed by our own ribosome profiling data. Thus, in terms of translational output ACO1/IRP1 shows >5-fold higher expression than IREB2/IRP2 (Fig. 2B; compare maximal liver footprint Reads Per Kilobase per Million mapped reads of ∼150 for *Aco1* and of ∼25 for *Ireb2*). Of note, however, previous studies that failed to show major phenotypes in IRE regulation upon IRP1 loss did not address the possibility that IRP1-mediated effects in liver might be subject to temporal control over the diurnal cycle.

We thus wished to determine experimentally how rhythmic target regulation would be affected in the absence of IRP1 or IRP2. To this end, we made use of *Aco1* and *Ireb2* knockout mouse strains [37]. These mice present with systemic alterations to iron homeostasis, including in liver (reviewed in [38]); moreover, there are documented links between iron and the circadian timekeeping mechanism itself. For example, several core clock proteins contain heme [39–41] and the transcription of *Per1* and *Per2* has been reported as sensitive to iron accumulation through a mechanism that involves histone methylation state [42]. A more systematic analysis of how iron metabolic genes affect the circadian clock is still lacking in mammals, but has been carried out in *Drosophila* and revealed numerous interactions [43]. Therefore, for our study we first explored whether whole-body *Aco1* or *Ireb2* knockout would engender phenotypes in entrained liver clock rhythms that could potentially confound the specific analysis of rhythmic IRP/IRE regulation. We used a method for the real-time recording of liver gene expression in freely moving mice [44, 45] that relies on luciferase reporters, luciferin delivery via an osmotic minipump, and highly sensitive bioluminescence photon counting (Fig. 3A). All animals carried the *mPer2::luc* knock-in allele [46] that expresses a protein fusion between core clock component PER2 and firefly luciferase, serving as a precise readout of diurnal oscillations. We carried out real-time recordings of liver rhythms under conditions that ensured light-entrainment of the master (SCN) clock to an external 24-hour light-dark cycle by means of a skeleton photoperiod, i.e. two 30 min light pulses applied at times corresponding to the beginning and to the end of the light phase in a 12h-light-12h-dark (LD12:12) cycle (Fig. 3B). Our experiments revealed rhythmic bioluminescence rhythms in wild-type animals, as expected, and also in littermate animals that were knockouts for either of the IRPs (Fig. 3C-F). For *Aco1^-/-^*mice, the phase of entrained mPER2::LUC reporter oscillations was virtually indistinguishable from controls (Fig. 3D). In the *Ireb2^-/-^* animals, the entrained mPER2::LUC phase was minimally delayed (∼0.5 h) as compared to controls (Fig. 3F), yet no other overt differences were apparent. We nevertheless verified that the liver clock in *Ireb2* knockouts was fully functional by recording free-running rhythms from liver organ explants (Fig. 3G). Free-running bioluminescence rhythms from *Ireb2* knockouts and controls (Fig. 3H) and their quantification (Fig. 3I) indicated overall comparable oscillations *ex vivo*, yet with higher variability in period length between individual explants in the knockouts. Altogether, our data suggested that diurnal oscillations in liver were comparable between IRP knockouts and their respective controls. Entrained clock phases between genotypes showed good correspondence.

**Figure 3.**
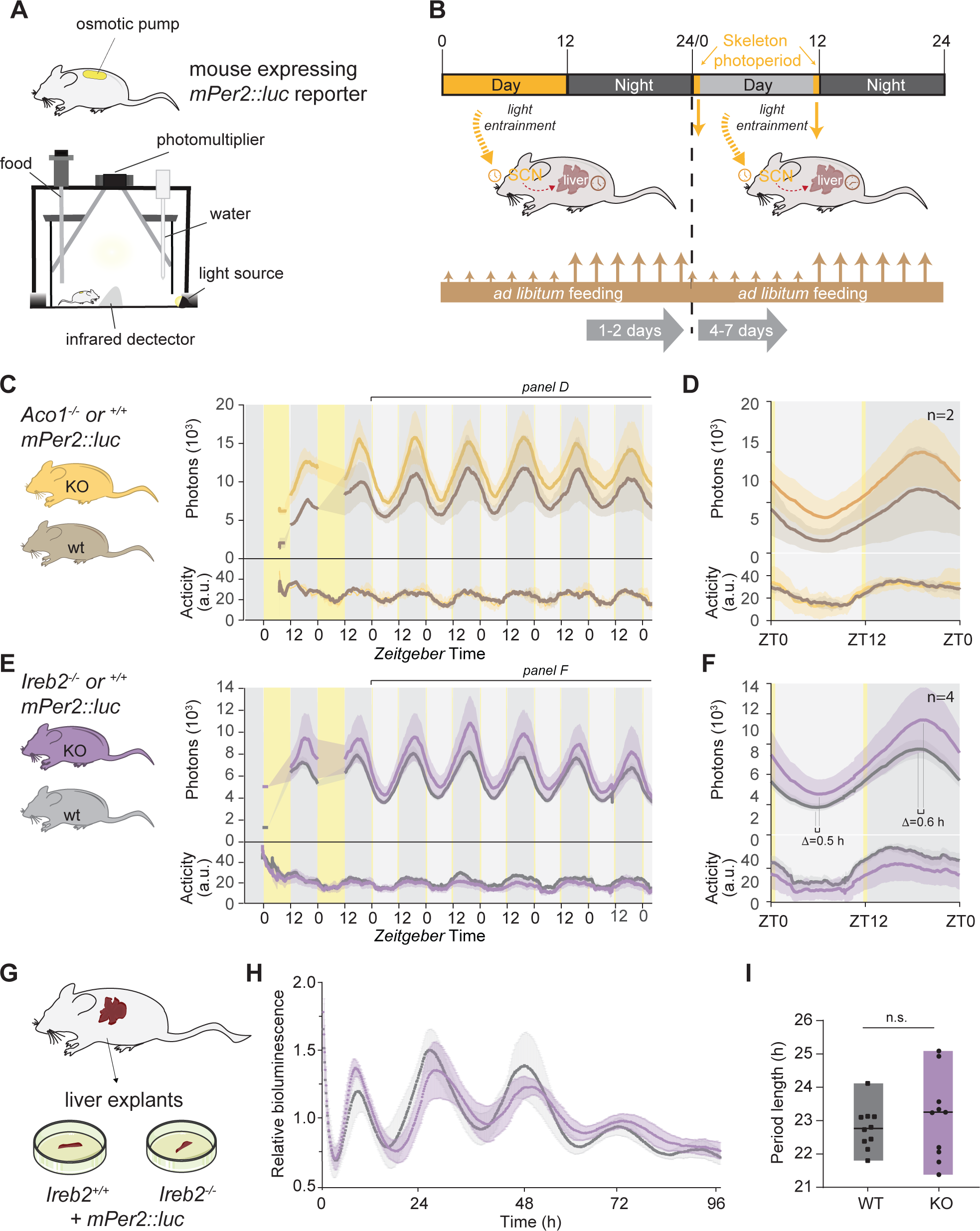
*In vivo* bioluminescence recoding shows that entrained circadian rhythms in liver are unperturbed in mice that are deficient for *Aco1* or *Ireb2*. (**A**) Cartoon depicting the *in vivo* recording setup (RT-Biolumicorder) used to measure diurnal bioluminescence rhythms in freely moving mice. The mouse is implanted with a luciferin-charged osmotic minipump and dorsally shaved to ensure detectability of photons emitted from the liver. It is then placed in the arena of the RT-Biolumicorder that records photons via a photomultiplier tube (PMT), activity via infrared detection, and also offers the possibility to program food availability and illumination. (**B**) Schematic representation of the applied recording protocol in the RT-Biolumicorder setup. Mice expressing the *Per2::Luc* reporter and implanted with the minipump are placed in the RT-Biolumicorder where they are first subject to LD12:12 schedule and *ad libitum* feeding for 1-2 days, allowing for acclimation to the setup. Bioluminescence recording only occurs during the dark phase. Then, the illumination schedule is changed to skeleton photoperiod, i.e. two 30 min light pulses applied at times corresponding to the beginning and to the end of the light phase in a 12h-light-12h-dark (LD12:12) cycle. Recording occurs throughout the day with the exception of the duration of the 30 min light pulses. The skeleton photoperiod is important as it keeps the SCN clock entrained to a defined 24-hour T-cycle, allowing to specifically query potential changes in the entrained liver clock. (**C**) Bioluminescence rhythms and activity traces recorded with the protocol shown in (B) from mice deficient for *Aco1* (ochre, N=2) and controls (grey, N=2). Mean signal (solid trace) and SEM (shaded) over the whole course of the experiment are shown. (**D**) Compiled data of (C), averaging all cycles from day 3. No change in PER2::LUC phase was detectable between genotypes. (**E**) Bioluminescence rhythms and activity traces recorded with the protocol shown in (B) from mice deficient for *Ireb2* (purple, N=4) and controls (grey, N=4). Mean signal (solid trace) and SEM (shaded) over the whole course of the experiment are shown. (**F**) Compiled data of (E), averaging all cycles from day 3. For *Ireb2^-/-^* mice, PER2::LUC peak and trough phases were slightly delayed as compared to the control mice with the same genetic background. (**G**) Schematic of liver explant rhythm measurements from the two genotypes, *Ireb2^-/-^* and controls, using male mice from the same breedings. Mice carried the *Per2::Luc* reporter allele. (**H**) Liver explant bioluminescence recording from *Ireb2^-/-^* (purple) and control (grey) mice. Free-running circadian traces show mean signal and SEM for the two genotypes over a recording period of 96 hours. N=10 per genotype. (I) Period length quantification of experiments shown in (H). No significant change in period length was detectable (Mann-Whitney test). *Ireb2* knockout period lengths appeared more variable than controls.

### Timepoint-specific gene expression datasets from IRP-deficient livers allow analysis of diurnal regulation

For our further analyses we selected the two timepoints at which rhythmic IRP/IRE activity would be low (ZT5) and high (ZT12), respectively, leading to high and low translation efficiency *(Ftl1, Fth1, Alas2)* and mRNA turnover *(Tfrc)* (Fig. 4A). We reasoned that in the simplest scenario, one of the IRPs (e.g. IRP2, the most potent IRE-regulator in liver according to the literature) was singly responsible for rhythmic target regulation. This rhythm would then be detectable in the control livers when comparing the two timepoints, yet absent in the livers from animals deficient for the rhythm-relevant IRP (e.g. IRP2 in the above example). More complex regulatory scenarios are conceivable, too, such as redundant regulation by the two IRPs, which would lead to no, or partial deregulation of rhythmicity in the knockouts. We collected 3 livers per timepoint and genotype (*Aco1^-/-^*and *Ireb2^-/-^*; plus their respective, matched *Aco1^+/+^*and *Ireb2^+/+^* controls carrying the same genetic background) and prepared RNA-seq and ribo-seq libraries (Fig. 4B). After sequencing and analysis, quantification of *Aco1-*mapping ribosome protected fragments (RPFs) in *Aco1^-/-^* livers confirmed the knockout at the level of ACO1/IRP1 translation/protein biosynthesis (Fig. 4C, top panel). Because in the knockout allele the open reading frame is disrupted yet still produces an mRNA from the locus [37], the reduction in *Aco1* in the RNA-seq data was less pronounced (Fig. 4C, middle panel). *Aco1* translation efficiency (TE; defined as RPF/RNA ratio) was decreased in the knockouts as expected (Fig. 4C, lower panel). In livers from *Ireb2*-deficient animals, *Aco1* RNA and RPFs were decreased by 10-20% as compared to their littermate controls (Fig. 4D), indicating that IREB2/IRP2 loss did not lead to compensatory higher IRP1 biosynthesis. Overall analogous observations were made for *Ireb2* expression, which showed the expected decrease in the *Ireb2^-/-^*animals at RPF and TE level (Fig. 4F, upper and lower panels), but not for the RNA (Fig. 4F, middle panel). In *Aco1* knockouts, *Ireb2* expression was largely unchanged at the RPF level, indicating that there was no substantial compensatory upregulation (Fig. 4E). We next validated knockout and possible compensatory effects by Western blot. These analyses confirmed the expected loss of ACO1/IRP1 in *Aco1^-/-^* mice (Fig. 4G, H). In these animals, IREB2/IRP2 abundance showed a tendency towards increased levels that did not reach statistical significance (Fig. 4G, I). Higher abundance of IREB2/IRP2 at ZT12 vs. ZT5 was evident as well, although quantitively less pronounced than in the around-the-clock time series analysed before (compare Fig. 4G, I with Fig. 2G-J). In the *Ireb2* cohort, knockouts were devoid of IREB2/IRP2 protein as expected (Fig. 4J, L). In the control animals from this cohort, IREB2/IRP2 increased in abundance from ZT5 to ZT12 in a quantitatively minor (yet statistically significant) fashion (Fig. 4L). In summary, we concluded that the datasets obtained from the two timepoints and genotypes would allow the analysis of interactions between IRP function and time-of-day and, more specifically, to investigate whether the identified rhythmic regulation could be attributed specifically to one of the IRPs. Moreover, our experiments validated the specificity of the IRP1 and IRP2 antibodies. Finally, although we observed a significant difference in IRP2 abundance between light-phase and dark-phase timepoints across different independent datasets, the fold-changes seen in the three cohorts (Fig. 2I, J; Fig. 4I, L) were highly variable, pointing towards poor robustness of IRP2 rhythmicity.

**Figure 4.**
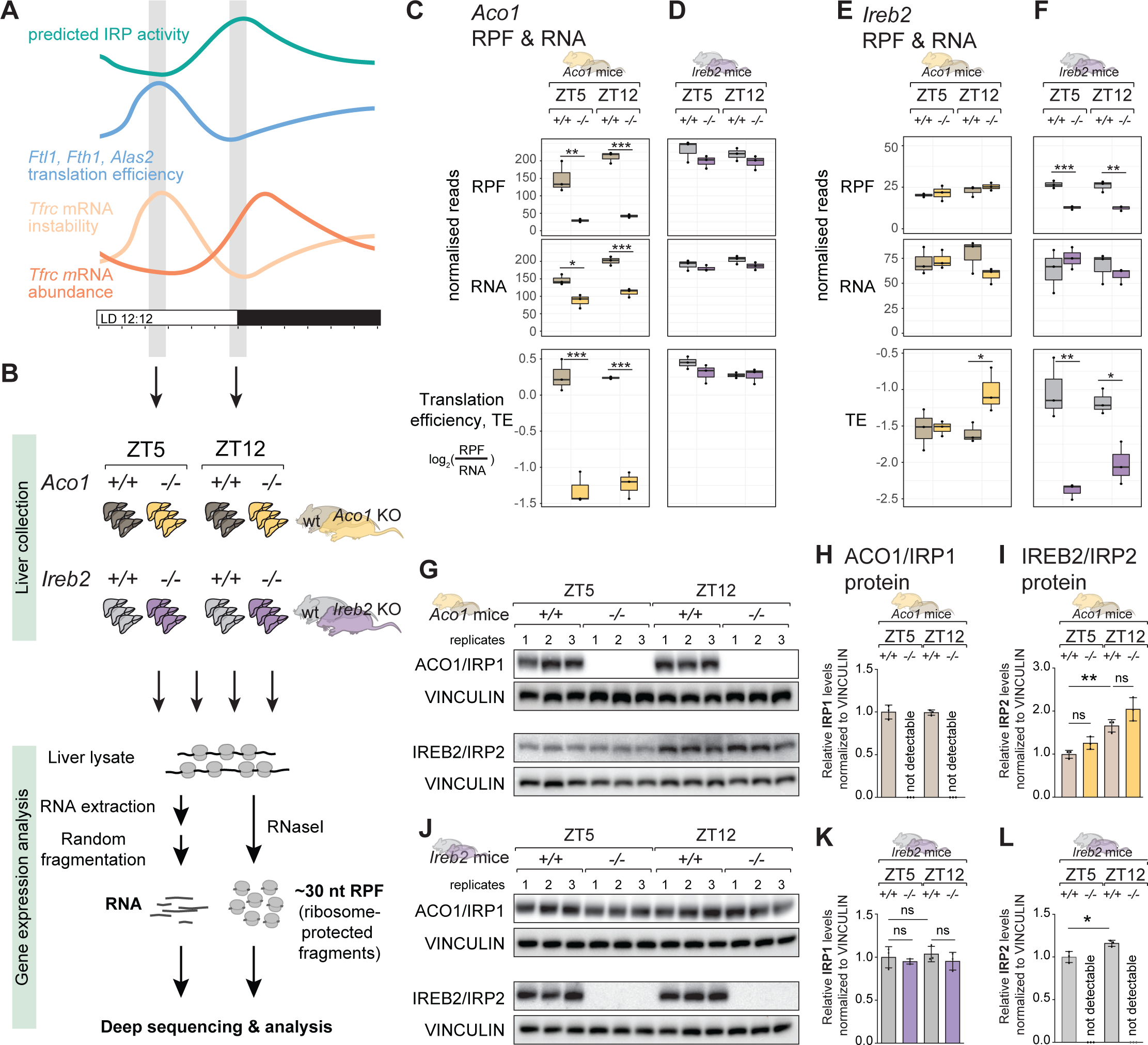
Time-resolved datasets allow analysis of IRP1 vs. IRP2-dependence of rhythmic regulation. (**A**) Schematic representation of the selection of timepoints for the ribo-seq experiment. Thus, high *Ftl1, Fth1* and *Alas2* translation efficiency and low *Tfrc* abundance, conceivably as a result of mRNA instability, occur in the light phase of the day, leading us to predict low IRP activity around ZT5. By contrast, low *Ftl1, Fth1* and *Alas2* translation efficiency and the rise in *Tfrc* abundance at the light-dark transition suggests high IRP activity at ZT12. (**B**) Schematic of the experiment carried out to assess timepoint- and genotype-specific ribosome profiling in the two IRP-deficient mouse strains and matched controls. Thus, 3 animals each were sacrificed at ZT5 and ZT12, for genotypes *Aco1^+/+^*(dark grey), *Aco1^-/-^* (ochre), *Ireb2^+/+^* (light grey), *Ireb2^-/-^* (purple). Livers were further processed for ribo-seq and RNA-seq. (**C**) Quantification of ribosome protected fragments (RPF, top) and RNA abundance (middle) and relative translation efficiency (log_2_ ratio of RPF/RNA) for the *Aco1* transcript in *Aco1* knockout mice and their matched controls, for the two timepoints ZT5 and ZT12. Read count is normalised to library depth. Statistical significance evaluated by unpaired t-test and indicated with asterisks (*** *P* < 0.001, ** *P* < 0.01, * *P* < 0.05). (**D**) As in (C), for the *Aco1* transcript in *Ireb2* knockout mice and their matched controls. (**E**) Quantification of ribosome protected fragments (RPF, top) and RNA abundance (middle) and relative translation efficiency (log_2_ ratio of RPF/RNA) for the *Ireb2* transcript in *Aco1* knockout mice and their matched controls, for the two timepoints ZT5 and ZT12. Read count is normalised to library depth. Statistical significance evaluated by unpaired t-test and indicated with asterisks (** *P* < 0.001, ** *P* < 0.01, * *P* < 0.05). (**F**) As in (E), for the *Ireb2* transcript in *Ireb2* knockout mice and their matched controls. (**G**) Western blot analysis of total protein extracts from the livers used for the gene expression profiling in panels (C) and (D), i.e. the *Aco1^-/-^* and matched *Aco1^+/+^* mice. VINCULIN was used as loading control to normalise ACO1/IRP1 and IREB2/IRP2 signals. (**H**) Quantification of Western blot signals for ACO1/IRP1 (normalised for loading via VINCULIN) from panel (G). No ACO1 is detectable in the *Aco1^-/-^* animals, as expected. ACO1 abundance is identical in the control animals at ZT5 and ZT12. (I) Quantification of Western blot signals for IREB2/IRP2 (normalised for loading via VINCULIN) from panel (G). IREB2 is slightly upregulated in the *Aco1^-/-^* vs. *Aco1^+/+^*animals (not significant). IREB2 levels are increased at ZT12 vs. ZT5 (** *P* < 0.01, One-way ANOVA with Šídák’s multiple comparisons test). (J) As in (G), but for the *Ireb2^-/-^* and matched *Ireb2^+/+^* animals whose livers were used for gene expression analysis in panels (E) and (F). (K) Quantification of Western blot signals for ACO1/IRP1 (normalised for loading via VINCULIN) from panel (J). Differences beween timepoints or genotypes (*Ireb2^-/-^* vs. *Ireb2^+/+^*) were statistically not significant (One-way ANOVA with Šídák’s multiple comparisons test). (L) Quantification of Western blot signals for IREB2/IRP2 (normalised for loading via VINCULIN) from panel (J). IREB2 is undetectable in *Ireb2^-/-^* mice as expected. The upregulation from ZT5 to ZT12 is statistically significant (* *P* < 0.05, Student’s t-test).

### IRP loss affects the regulation of 5’ and 3’ IRE-containing transcripts in a timepoint-specific fashion

Next, we assessed across genotypes and timepoints how 5’ IRE-containing transcripts (*Ftl1*, *Fth1*, *Alas2*) were affected in their translation efficiencies (Fig. 5A-C) and 3’ IRE-containing *Tfrc* in its RNA abundance (Fig. 5D). In the TE analyses, we observed changes that were not only IRP-specific, but also strongly dependent on timepoint. The most striking observation was the timepoint-specificity of translational derepression in *Ireb2^-/-^* mice (Fig. 5A-C, right panels). Thus, *Ftl1* and *Fth1* (and to a lesser extent *Alas2*) showed strongly increased TEs (i.e., derepression) in the absence of IREB2/IRP2 during the light phase of the day (ZT5), in line with the model that IREB2/IRP2 is the main IRE-regulating IRP in liver [14–16]. By contrast, at the beginning of the dark phase (ZT12), we observed that TEs were low and near-identical between *Ireb2^-/-^* and *Ireb2^+/+^* animals (Fig. 5A-C, right panels). These findings indicated that a repressive mechanism distinct from IREB2/IRP2 was acting on the translation of IRE-containing transcripts specifically at ZT12. This mechanism was unable to compensate for IREB2/IRP2 loss at the light phase timepoint. Timepoint-specific effects in *Ireb2^-/-^*vs. *Ireb2^+/+^* animals could also be observed for *Tfrc* (Fig. 5D, right panels). On the one hand, the tendency to reduced *Tfrc* levels in *Ireb2^-/-^* livers (in line with the model that IRP2 protects *Tfrc* mRNA from degradation) was visible at ZT5. However, at ZT12 there was no decrease in *Tfrc* mRNA in the absence of IREB2/IRP2, and rather a trend towards increased abundance. Therefore, similarly to the TE repressive effects seen for 5’ IREs, also the mRNA stabilizing effect for 3’ IRE-containing *Tfrc* did not require IRP2 at the dark-phase timepoint.

**Figure 5.**
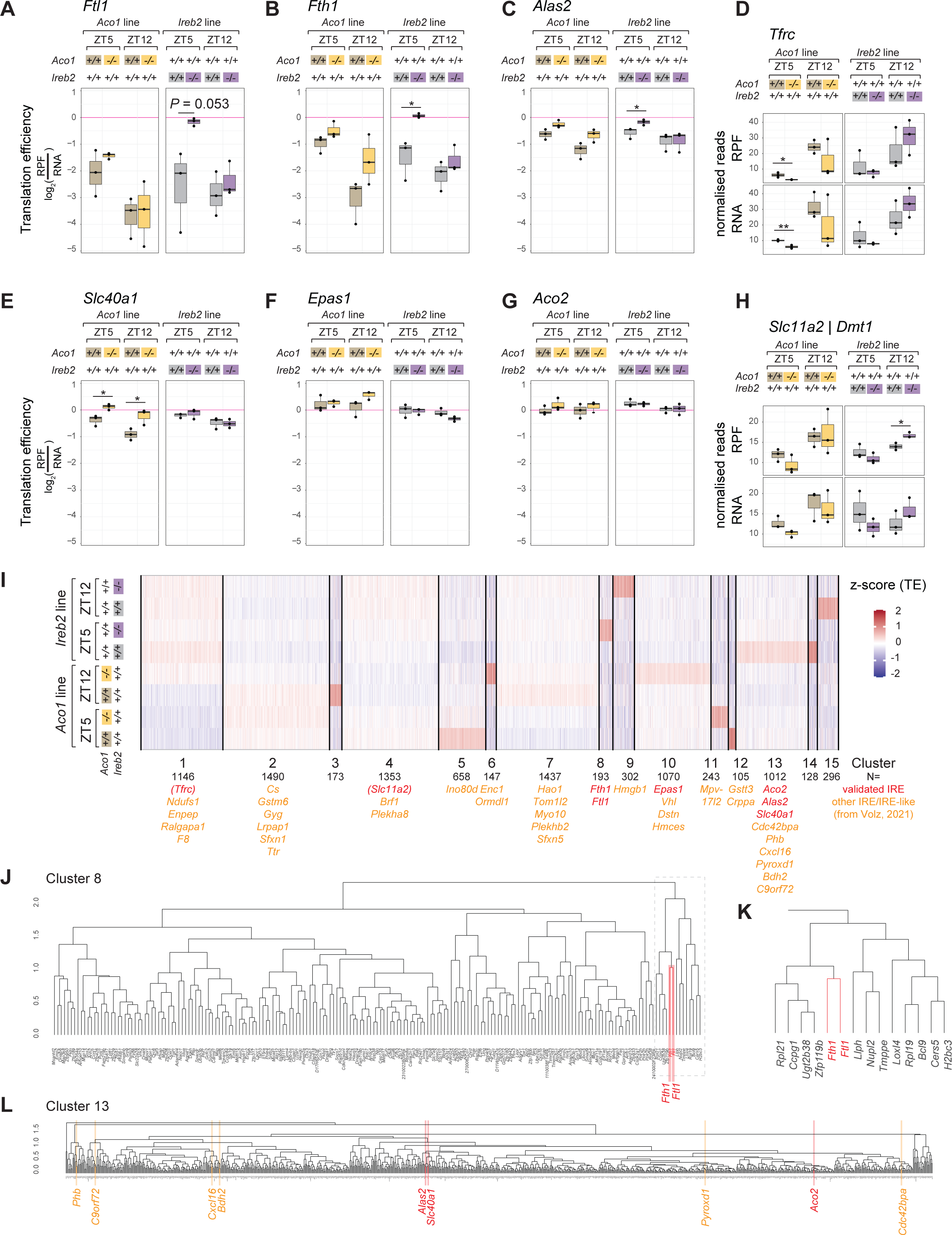
Timepoint- and transcript-dependence of derepression upon loss of ACO1/IRP1 vs. IREB2/IRP2. (**A**) Translation efficiency analysis for *Ftl1* at ZT5 and ZT12 in the *Aco1* line (*Aco1^-/^*^-^ vs. *Aco1^+/+^*in ochre and grey, respectively) in the left panel and for *Ireb2* line (*Ireb2^-/-^* vs. *Ireb2^+/+^* in purple and grey, respectively) in the right panel. Statistical significance evaluated by unpaired t-test and indicated with asterisks (*** *P* < 0.001, ** *P* < 0.01, * *P* < 0.05). (**B**) As in (A), for *Fth1*. (**C**) As in (A), for *Alas2*. (**D**) RPF (upper) and RNA (lower) normalised reads for *Tfrc* at ZT5 and ZT12 in the *Aco1* line (*Aco1^-/^*^-^ vs. *Aco1^+/+^*in ochre and grey, respectively) in the left panel and for *Ireb2* line (*Ireb2^-/-^* vs. *Ireb2^+/+^* in purple and grey, respectively) in the right panel. Statistical significance evaluated by unpaired t-test and indicated with asterisks (*** *P* < 0.001, ** *P* < 0.01, * *P* < 0.05). (**E**) As in (A), for *Slc40a1*. (**F**) As in (A), for *Epas1*. (**G**) As in (A), for *Aco2*. (**H**) As in (D) for *Slc11a2/Dmt1*. (I) K-means clustering analysis on z-score-transformed TE profiles across all eight conditions (2 timepoints, 4 genotypes; replicate means were used). The numbering of the 15 clusters and the gene count are given below the heatmap. Below the clusters, the known/validated IRE-containing transcripts are highlighted in red, with (*Tfrc* and *Slc11a2* in parantheses, given that their IRE-regulation is not at the level of TE, but of RNA stability). Predicted IRE-like mRNAs from a list in [47] are highlighted in orange. Note that the two most rhythmically translated IRE-containing mRNAs, *Ftl1* and *Fth1* are found together in cluster 8. Several other IRE transcipts are enriched in cluster 13. (J) Hierarchical clustering according to the similarity in TE profiles for transcripts in cluster 8 from panel (I). *Ftl1* and *Fth1* cluster together (marked in red). (K) Zoom into the part of cluster 8 that contains *Ftl1* and *Fth1* in direct vicinity (marked in red). The unsupervised clustering thus confirms that these two transcript indeed have a distinct mode of (rhythmic) regulation that sets them apart from other transcripts, even IRE-containing ones. (L) Hierarchical clustering according to the similarity in TE profiles for transcripts belonging to cluster 13 from panel (I), which contains several validated (red) and predicted (orange) IRE-containing transcripts, possibly indicating common regulatory mechanisms and, in particular, similar “set-points”.

We next analysed the effects associated with the absence of IRP1 (*Aco1^-/-^* vs. *Aco1^+/+^* animals). Previous work has demonstrated that FTL1 protein levels are unchanged in livers of IRP1-deficient mice, including under low- and high-iron diets [14]. Indeed, *Ftl1* TE was overall similar between *Aco1^-/-^*and *Aco1^+/+^* animals (Fig. 5A, left panel). For *Fth1* and *Alas2,* some translational derepression was visible at ZT5 and, in particular for *Fth1*, also at ZT12 (Fig. 5B, C, left panels; effects not statistically significant). As a result, the differences between TEs at peak (ZT5) and trough (ZT12) were reduced in *Aco1^-/-^*vs. *Aco1^+/+^* animals, indicating slightly blunted amplitudes of rhythmic *Fth1* and *Alas2* translation when ACO1/IRP1 was absent. Finally, changes in *Tfrc* mRNA abundance were overall consistent with the effects seen for *Fth1* and *Alas2* TEs. Thus, *Tfrc* mRNA levels were lower in the absence of IRP1 at both timepoints (Fig. 5D, left panels; statistically significant at ZT5 but not at ZT12 due to variability across biological replicates). We concluded that despite the generally held model that IRP1 has little effect on IRE regulation in liver, specific differences between knockouts and controls were apparent. However, these did not reach statistical significance due to the relatively low sample size (3 animals per group) and considerable variability across mice.

In summary, the above analyses uncovered a complex relationship between timepoint (ZT5, ZT12) and genotype (*Aco1^-/-^* vs. *Aco1^+/+^*; *Ireb2^-/-^* vs. *Ireb2^+/+^*) in the regulation of several IRE-containing mRNAs. Our data indicate that IRP2 is not solely responsible for IRE regulation in liver. Especially at ZT12, IRP2 actually appears to be dispensable for translational repression. Our analyses further revealed that absence of IRP1 affects the regulation of IRE-containing mRNAs, in particular at ZT12. Taken together, it is conceivable that for rhythmically regulated IRE transcripts, IRP activities are partitioned and partially redundant according to time-of-day, with IRP2 dominating regulation at ZT5 (light phase) and IRP1 able to act at ZT12 (dark phase).

### Distinct timepoint-dependent regulation and basal repressed states distinguish core iron metabolic and other 5’ IRE-containing transcripts

Beyond the four transcripts analysed in the previous section, several other mRNAs contain validated IREs (e.g. Fig. 1D), and many more have been predicted [47]. A recent study has proposed that across different IRE-containing transcripts in rat liver, translational control through IRPs follows a hierarchy based on different iron concentration set-points at which translational repression occurs [48]. Thus, IRE-containing mRNAs encoding core iron storage factors (e.g. Ferritins) are translationally repressed under normal physiological conditions and chow diet (i.e., “regular” cellular iron levels; our animals were fed normal chow diet throughout), whereas those mRNAs that encode proteins with broader physiological roles are actively translated under these conditions (e.g. *Aco2*, encoding the TCA cycle enzyme, mitochondrial aconitase). Under iron overload, the former transcripts would undergo derepression, yet the latter only experience little additional stimulation of translation. Upon iron deficiency, by contrast, *Aco2* is highly responsive and becomes strongly repressed. Interestingly, it was proposed that IRP1 contributes to this setpoint mechanism through the kinetics of IRP1:IRE dissociation [48]. Conceivably, similar mechanisms may also underlie the differences that we observe in rhythmic regulation. Indeed, in the control mice, *Ftl1* and *Fth1* (encoding core iron management proteins) were generally more strongly repressed than *Alas2* (encoding a broader function protein involved in heme biosynthesis) – see TE values in the wild-type animals in Fig. 5A-C that were in the range of −4 to −1 for *Ftl1* and *Fth1*, but around −1.5 to −0.5 for *Alas2*. Note that TE values are derived from RPF/RNA ratios and can be compared across different transcripts quantified from the same libraries.

In order to investigate this phenomenon more globally, we set out to analyze the TEs of other IRE-containing mRNAs in our datasets. First, we explored the group of transcripts with documented 5’ IREs that only presented weak, or no, rhythmicity in our original dataset [6] and that are shown in Fig. 1D. Consistent with the validity of the “setpoint model”, these transcripts all showed high basal TEs (−1 to +0.5) and IRP deficiency led, at most, to modest stimulation of translation (Fig. 5E-G). Thus, in *Ireb2^-/-^* animals, *Slc40a1/Ferroportin*, *Epas2/Hif2a* and *Aco2* were only weakly altered in their TEs at either timepoint (Fig. 5E-G, right panels). In *Aco1/Irp1* knockouts, significant derepression was observable at both ZT5 and ZT12 for *Slc40a1/Ferroportin* (encoding a protein in the core iron metabolic machinery that functions as an iron exporter [48]; Fig. 5E, left panel). For *Epas1/Hif2a* a small effect was apparent at ZT12 (Fig. 5F, left panel; statistically not significant; please note that a previous report had suggested specific *Epas1/Hif2a* regulation through IRP1 in liver [16]). The effect size of de-repression for *Aco2* was quantitatively minimal as well (Fig. 5G, left panel). Finally, we also analysed possible gene expression changes for *Slc11a2/Dmt1*, which bears an IRE in its 3’ UTR [49]. *Slc11a2* RNA levels changed in a similar fashion to what we had observed for *Tfrc* upon loss of IRP1 or IRP2 (compare Fig. 5H to Fig. 5D).

Next, we set out to classify transcripts in a comprehensive fashion, according to how their translation rates were affected across conditions in our ribo-seq datasets. To this end, we used k-means clustering to partition the transcriptome into overall 15 clusters according to the similarity of their z-score-transformed TE profiles (Fig. 5I and Table S1). Validated IRE-containing transcripts and predicted IRE-like mRNAs (list from [47]) were found in several clusters. The two most rhythmically translated IRE-containing mRNAs, *Ftl1* and *Fth1* – but no other (validated or predicted) IRE transcripts – were found together in a common cluster (cluster 8), where they were grouped in immediate neighbourhood (Fig. 5J, K), consistent with their distinct strongly rhythmic regulation. Another cluster (cluster 13) contained several 5’ IRE transcripts (*Aco2, Alas2, Slc40a1*, and predicted IRE-like candidates; see Fig. 5I, L), in line with common regulatory mechanisms and possible similar “set-points”. Within cluster 13, the close vicinity of *Alas2* and *Slc40a1*, as compared to the more distant *Aco2* (Fig. 5L), recapitulated the differences in magnitude of derepression seen in the individual transcript analyses shown in Fig. 5C, E, G.

In summary, the above analyses support the idea that the observability of rhythmicity on IRE-containing transcripts depends on whether in the basal state, these mRNAs are strongly repressed, which allows for a timepoint-dependent derepression as a function of reduced IRP activity.

### IRP/IRE rhythmicity follows feeding-fasting cycles and correlates with hepatic ferrous iron measurements

We next aimed to gain insights into upstream regulatory mechanisms that drove the observed IRP/IRE rhythms. First, we asked the question if the phenomenon was *bona fide* circadian clock-driven that depended on functional clocks locally in hepatocytes and/or systemically in the whole organism, or if other rhythmicity cues were involved. Briefly, it is well established that various direct and indirect mechanisms, alone or in combination, are involved in driving gene expression oscillations in liver (reviewed in [50]). A considerable proportion of rhythmic mRNA expression is thus driven transcriptionally and directly through the cellular core clock. However, numerous cases of hepatic gene expression oscillations still persist in the absence of local liver clocks, as they are driven by systemic cues. The underlying mechanisms include, for example, oscillations in body temperature that are generated by the master clock in the SCN and lead to changes in the activity of the transcription factor *Hsf1*, which in turn drives transcriptional oscillations without the obligatory involvement of local hepatocyte clocks [51–53]. Moreover, feeding/fasting cycles (that are a consequence of behavioural rhythmicity) are an important timing cue for metabolically active peripheral organs. Rhythmic feeding is a well-established *Zeitgeber* for the entrainment of hepatic clocks and, in addition, feeding-dependent signals also regulate gene expression directly, without the need of local hepatocyte clocks. For example, the nutrient-responsive mTOR pathway undergoes daily activation cycles in response to feeding, regulating the translation of ribosomal protein mRNAs containing regulatory 5’ TOP motifs [6, 54, 55], showing maximal translation at the light-dark transition, i.e. in antiphase to IRP/IRE translation rhythms.

In order to explore at which level in the organism a circadian clock was required for IRP/IRE rhythms, we used the oscillations of *Tfrc* mRNA as a proxy for rhythmic IRP/IRE regulation. This allowed us to take advantage of numerous published around-the-clock RNA-seq datasets from wild-type and clock-deficient mice under a variety of feeding regimes. First, we analysed the RNA-seq data from a recent study [56] that used three different genetic animal models: wild-type, *Bmal1* knockout (i.e., behaviourally arrhythmic with no functional clocks in the entire body) and liver-specific *Bmal1* re-expression (i.e. behaviourally arrhythmic, with functional clocks only in hepatocytes), either fed *ad libitum* or night-restricted. Night-restricted feeding exacerbated *Tfrc* rhythmicity in wild-type mice (Fig. 6A, upper). In *Bmal1* KO mice, *Tfrc* was arrhythmic under *ad libitum* feeding, yet rhythmicity was re-installed under night-feeding (Fig. 6A, middle) and in *Bmal1* re-expression mice, *Tfrc* regained rhythmicity under night-feeding (Fig. 6A, lower). This data was fully compatible with a model according to which local clocks (liver) or a circadian master clock (SCN) were *per se* not essential for *Tfrc* rhythmicity, as long as feeding-fasting cycles persisted. We confirmed this conclusion in two other datasets. Atger et al. [7] collected livers from night-fed full-body *Bmal1* KO vs. wild-type mice and applied an RNA-seq methodology to quantify both mRNA and pre-mRNA levels. In this experiment, *Tfrc* mRNA rhythms are indistinguishable between the circadian clock-deficient *Bmal1* KO and the control animals under imposed night-feeding (Fig. 6B, upper panel), and pre-mRNA (transcription) is identical and arrhythmic in both genotypes (Fig. 6B, lower panel). We concluded that rhythmic feeding was sufficient to induce post-transcriptionally generated *Tfrc* mRNA oscillations in the absence of an endogenous circadian system. Next, we used the study by Greenwell et al. [57] to investigate the complementary scenario, i.e. how *Tfrc* rhythmicity in liver fares in the presence of a wild-type circadian system, but in the absence of feeding/fasting cycles (Fig. 6C). The analysis of four different feeding regimens from night-restricted (100% food intake during the dark phase) to *ad libitum* (∼76% during dark), “damped”/almost entirely arrhythmic feeding (∼56% during dark) and arrhythmic feeding (∼51% of food intake during dark vs. ∼49% during light phase), revealed a gradual disappearance of *Tfrc* rhythmicity that scaled with the loss of feeding/fasting cycles (Fig. 6C). Finally, we analysed data comprised in the study by Manella et al. [58] in which wild-type mice were subjected to *ad libitum* or to light phase-restricted feeding (using an experimental protocol known to have little influence on the phase of the liver clock), which revealed that the peak in *Tfrc* mRNA accumulation was inverted and followed feeding rhythms rather than the light-dark cycle (Fig. 6D). From the above analyses that probe the relative importance of the circadian system vs. feeding/fasting cycles for the rhythmic accumulation of *Tfrc*, we can conclude that it is via the latter mechanism – feeding – that *Tfrc* mRNA oscillations are engendered. By extension, we propose that feeding-related timing cues constitute a physiological basis for rhythmicity of the IRP/IRE system in liver tissue.

**Figure 6.**
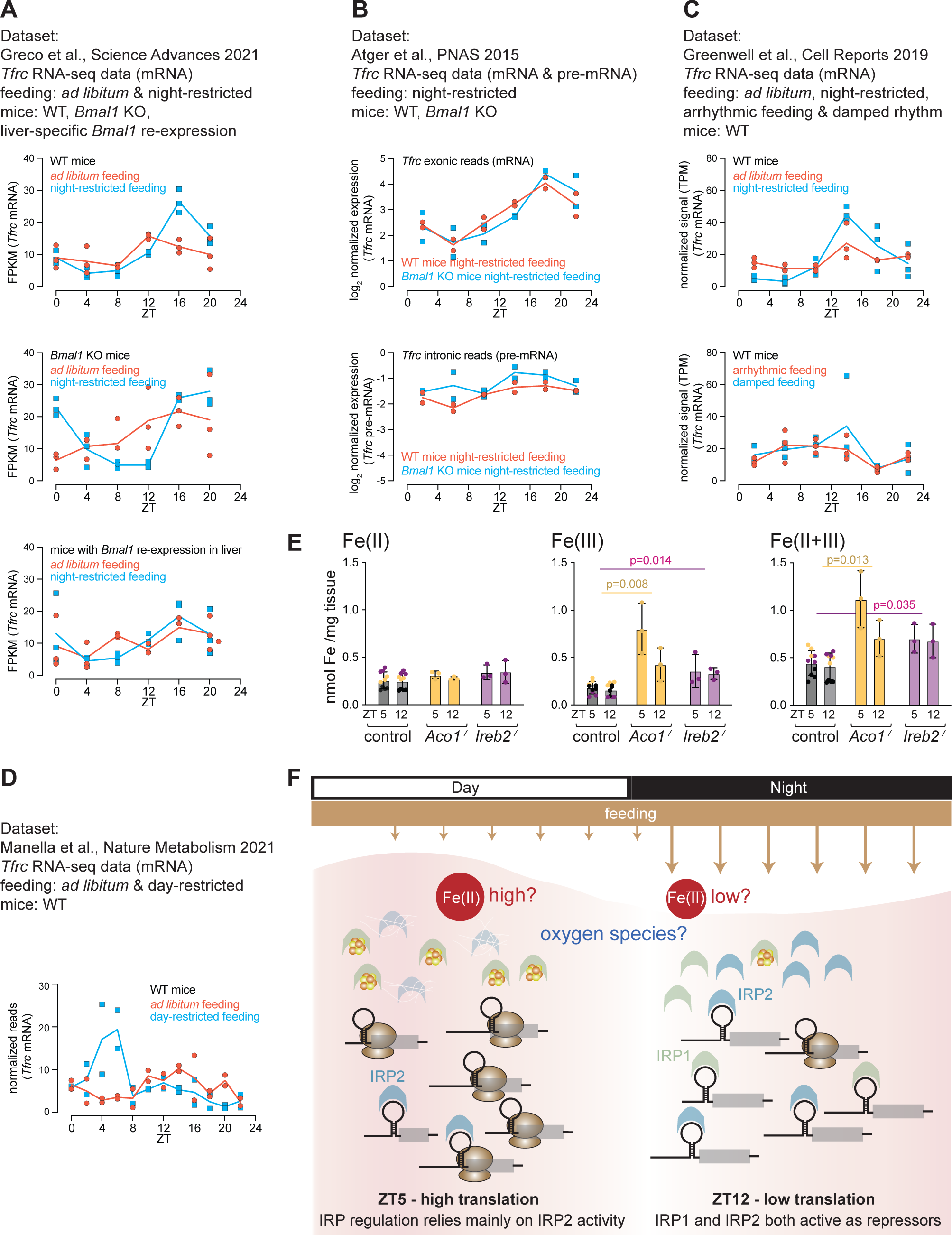
*Tfrc* oscillations follow feeding/fasting rhythms rather than the circadian clock. (**A**) Plotting of liver *Tfrc* expression data from the study by Greco et al. (source: Data S1) [56], which included two feeding paradigms (*ad libitum* vs. night-restricted) and three genotypes (WT, full-body clock-deficient *Arntl/Bmal1* KO, liver re-expression *Arntl/Bmal1*). The upper panel shows that night-restricted feeding exacerbates rhythmicity. The middle panel shows that night-restricted feeding restores rhythmicity in an otherwise circadian clock-deficient animal. A similar outcome is seen in the *Bmal1* re-expression animals shown in the lower panel. (**B**) Analogous to (A) using the data from Atger et al. (source: Dataset S1) [7] that contains both mRNA and pre-mRNA reads. Thus, under night-restricted feeding, wild-type and clock-deficient *Arntl/Bmal1* knockout animals show near-identical *Tfrc* oscillations (upper panel). These oscillations are post-transcriptionally generated, given that *Tfrc* pre-mRNA levels are not rhythmic (lower panel). (**C**) Analogous to (A) and (B) using the data from Greenwell et al. (source: Table S2) [57]. The upper panel compares *ad libitum* and night-restricted feeding effect on *Tfrc*, with the latter showing higher amplitude rhythms. The lower panel shows the loss of *Tfrc* rhythms in wild-type animals (i.e., containing a functional clock) when feeding becomes gradually less rhythmic. (**D**) Analysis of dataset from Manella et al. (source: Supplementary Data 2) [58] which compares wild-type animals that are *ad libitum*-fed vs. exclusively day-fed. Day-feeding inverts the *Tfrc* rhythmicity. (**E**) Measurement of tissue iron levels at ZT5 and ZT12 across liver samples from the *Aco1* KO/controls in ochre and the *Ireb2* KO/controls in violet (same animals as in Figure 4). 8 additional liver samples from independent, age-matched male mice (C57BL/6 background) were included, in black, to obtain more replicates in the controls. Still, differences between ZT5 and ZT12 were not significant. Indicated comparisons between *Aco1* KO vs. controls and *Ireb2* KO vs. control were significant for Fe(III) and Fe (II+III), in line with iron storage phenotype. Two-tailed, unpaired, parametric t-test. (**F**) Proposed model for rhythmic regulation of IRP/IRE activity. Left side - during the light phase (ZT5, coinciding with fasting phase in *ad libitum* fed mice): IRP2 levels are relatively lower, but still able to exert some translational inhibition on their targets. This is why in *Ireb2/Irp2*-deficient animals, depression is seen. IRP1 is less active, likely due to Fe-S cluster assembly that precludes its IRE binding and this may be brought about by increased Fe(II) levels or other cues such as oxygen flucations. Right side – at beginning of dark phase (ZT12, feeding phase): IRP2 is abundant and actively repressing, but IRP1 is also active and can compensate for the loss of IRP2 in the corresponding *Ireb2/Irp2*-deficient animals.

Iron itself controls IRE-binding activity for IRP1 and protein stability for IRP2 (Fig. 1B). Rhythmicity in intracellular, bioactive iron (the so-called “labile iron pool”, or LIP [59]) could therefore represent a plausible driver for the observed diurnal gene expression changes. However, the measurement of the relevant levels of Fe(II) – i.e., ferrous iron that is available to engage in regulation on IRPs in the cytoplasm of the hepatocytes, rather than more stably bound in other Fe-S-cluster containing proteins – next to other Fe iron pools and sources within the tissue, is a major challenge in the field (see Discussion). To gain insights into possible daily variations in liver iron, we adapted a method allowing the quantification of total, ferrous and ferric iron and that is insensitive to abundant heme-iron (see Materials and Methods). Our analyses revealed no significant changes in Fe(II) levels across the genotypes analysed (Fig. 6F). Given the methodological uncertainty surrounding our attempts to quantify the LIP in these experiments, we refrain from drawing strong conclusions on whether diurnal IRP/IRE regulation involves fluctuations in iron levels or alternative cues (see Discussion).

## Discussion

Through decades of research, the IRP/IRE system has become a paradigm (and literally a “textbook example” [60]) of post-transcriptional gene regulatory control that is, moreover, relevant for human diseases such as iron storage disorders [38]. The regulatory parameter that we have explored in our study – the time-of-day – has so far gone largely unnoticed (with the exception of a recent preprint [28]). We find high-amplitude rhythmic regulation in the liver precisely for the subset of transcripts that are archetypes of IRE regulation, namely the Ferritins (*Fth1, Fth2*), *Alas2* and *Tfrc*, which were also historically the first to be discovered [8–10, 24]. Previous analyses in IRP knockout livers have led to the model that IREB2/IRP2 is the main actor in hepatic IRE regulation and that ACO1/IRP1 cannot compensate for its loss (e.g. [14–16, 37]). None of these studies, however, used ribo-seq to specifically quantify translation rates, nor did they consider the temporal component in regulation. The timing of animal sacrifice was not reported in previous publications on hepatic IRE regulation, yet it is likely that it occurred during “core working hours” of laboratory researchers, i.e. between early morning and late afternoon. In particular the timepoint for which we detect the strongest IRP activity on its rhythmic targets and a possible redundancy between IRP1 and IRP2 activities – ZT12, at the beginning of the dark phase – is unlikely to have been covered in previously published work. Dark phase timepoints are, however, particularly relevant when working with nocturnal species, as this is when mice are most active and ingest most of their food.

By leveraging publicly available datasets and using *Tfrc* oscillations as a readout for diurnal IRP/IRE regulation, we uncovered that rhythmic control does not require a local circadian clock in hepatocytes, nor a central pacemaker in the SCN, as long as feeding/fasting cycles are maintained (Fig. 6). Daytime-fed mice have inverted hepatic *Tfrc* rhythms further underscores the dominant role of feeding (Fig. 6D). Hence, the direct influence and interference from the circadian clock onto the observed rhythm seems to be rather limited. In this aspect, IRP/IRE rhythmicity distinguishes itself from the mechanism acting on ribosomal protein (RP) mRNAs whose translation is also rhythmic and dependent on feeding. RP mRNA translation is increased at the light-dark transition in *ad libitum*- or night-fed mice [6, 7, 54]. However, in day-fed animals, the rhythm in newly synthesised RPs does not invert but becomes flat [55]. RP mRNAs contain 5’ TOP motifs that are regulated via the mTOR pathway, which shows multiple interactions with clock regulation [61]. While the translational upregulation of RP mRNAs by feeding is hence gated in a circadian fashion (i.e. the feeding signal can exert its stimulatory effect only at a specific time of day), no such circadian gating appears to be in place for the IRP/IRE system, as judged by *Tfrc* mRNA profiles.

Over the years, IREs have been reported, validated, or predicted in many transcripts – yet for none of them have we observed comparably high peak-to-trough amplitudes across the various datasets that we have analysed. In line with the “set-point model” that has been put forward to explain how IRP activity thresholds lead to a hierarchy of translational repression across 5’ IRE-containing transcripts [48], the most rhythmically regulated mRNAs are also the most sensitive to repression and generally exhibit lower translation rates (Fig. 5A-C) compared with the non- (or less-) rhythmically regulated transcripts, which are overall more highly translated (Fig. 5E-G). Therefore, we deem it likely that the appearance of rhythmicity on specific IRE-containing transcripts is related to whether the transcript/IRE-specific thresholds match the physiological dynamic range of regulatory cues that modulate the IRPs. This model would also explain why rhythmic regulation for the same mRNAs is less prominent in the kidney; conceivably, the dynamic range of the cues could be different in this organ.

A question that we have begun to address but that will require future investigations, concerns the identity of the molecular cue that is responsible for imparting rhythmicity. The prime candidate is certainly iron itself, more particularly iron in the so-called “labile iron pool” or LIP, which consists of redox-active iron ions that are weakly liganded and therefore engage in regulatory and metabolic processes [59]. Quantifying the relevant iron pool in tissues is technically not a facile enterprise, given the large amounts of iron that is stored in ferritin or otherwise bound in various Fe-S cluster-containing proteins, plus the large quantities of iron in the mitochondria and lysosomes. Our measurements show minimally higher Fe(II) at ZT5 than ZT12 (non-significant), i.e. in principle the phase that would be required for high and low IRP activity at ZT5 and ZT12, respectively. One concern that remains is that technical reasons preclude quantification of the relevant cytoplasmically active iron pool against large amounts of protein-bound Fe(II) that the assay we used measures as well; more suitable methodology for LIP measurements in animals would be an important advance in the field. Moreover, it is conceivable that additional cues impart rhythmicity on IRPs as well, in particular oxygen, reactive oxygen species as well as nitric oxide [36, 62–64]. Interestingly, daily rhythms in tissue O_2_ [65] and H_2_O_2_ [66] levels have been described. It will thus be important, yet not trivial, to disentangle to what extent these various cues are relevant for rhythmic regulation of IRE-containing transcripts.

Our study allows refining the role of IRP2 in hepatic IRE regulation and a possible interplay with IRP1. While there is indeed a large body of literature supporting the idea that IRP2 is the principle actor in the regulation of Ferritin protein and *Tfrc* mRNA accumulation in liver (e.g. [14–16]), the vast majority of former studies likely investigated regulation during the light phase (as lined out above). Our examination of time-resolved ribo-seq data indeed confirms that at the light phase timepoint ZT5, knockouts of IRP2 (but not of IRP1) present strong translational depression of Ferritin mRNAs. However, at the dark phase timepoint, ZT12, low translation efficiency is fully maintained even in the absence of IRP2 (Fig. 5A-B), indicating that a different repressor is capable of acting on these IREs at this time-of-day. An obvious candidate is IRP1, whose knockout indeed showed a tendency to derepression at ZT12, at least for *Fth1* (Fig. 5B). Since the double deficiency for both IRPs is not viable – even as a liver-specific knockout [67] – we cannot test the hypothesis that at ZT12, it is the combined activity of IRP1 and IRP2 that acts in repression. Still, based on our data we would propose a model (Fig. 6F) according to which IRP1 is inactive at ZT5, and thus all repression occurs through IRP2 for the light-phase timepoint, yet IRP1 and IRP2 are redundantly active at ZT12. In terms of possible molecular mechanisms that could engender higher IRP1 activity at this time point, lower iron levels or changes in one of the other relevant cues could play a role. For example, it is interesting to note that liver H_2_O_2_ levels (a reflection of overall mitochondrial activity) are highest precisely at the light-dark transition [66]. One may thus hypothesize that high H_2_O_2_ could engender the oxidation and destabilization of IRP1’s Fe-S cluster, leading to apo-IRP1 that is capable of binding to IREs. Experimental evidence will be needed to validate such speculations.

## Conclusions

Our investigations have brought to light a crucial aspect of regulation in the IRP/IRE system, namely its significant dependency on time-of-day. Through further analysis of this temporal dimension, we have uncovered previously overlooked redundant IRP activities notably at dark onset, a period likely overlooked in prior studies, thus challenging the notion of strict IRP2-dependence in liver. Our analyses further reveal that the daily fluctuations in IRP activity correlate with feeding-fasting cycles rather than with circadian cycles, and possibly involve hepatic Fe(II) level variations. Our findings thus reinforce the concept that rhythmic gene expression is governed by at least two distinct mechanisms. Firstly, truly circadian clock-driven oscillations, rooted in transcriptional regulation and essentially encoded within DNA sequence. Second, post-transcriptional/translational oscillations, frequently reflecting acute cues and signals such as feeding patterns. Translation emerges as a pivotal stage where such information can be ideally integrated to modulate protein output in a rapid and efficient manner. The responsiveness of the IRP/IRE system to feeding/fasting rhythms serves as a compelling example of this dynamic mechanism.

## Materials and methods

### Animals

Healthy adult male mice of age 12-24 weeks were used. All mouse lines were maintained on a C57BL/6J background. The knockout mice for IRP1 (allele *Aco1^tm1.1Mwh^*) and IRP2 (allele *Ireb2^tm1.2Mwh^*) have been described in [68]. Strains kindly provided by Matthias W. Hentze, EMBL Heidelberg.

### Generation of antibodies

Rat monoclonal antibodies against ACO1/IRP1 and IREB2/IRP2 were generated at the DKFZ Core Facility Antibodies. Briefly, full-length murine ACO1/IRP1 and IREB2/IRP2 proteins, fused to a poly-histidine tag, were expressed in *E. coli* and purified on Ni-NTA columns using standard protocols. Purified His-tagged proteins were used to immunise rats and generate hybridomas. Hybridoma supernatants were first screened by ELISA against His-tagged ACO1/IRP1 and His-tagged IREB2/IRP2. As an additional control, supernatants were tested against full-length His-tagged murine ACO2 (mitochondrial aconitase), which shares 27 and 26% identity with ACO1/IRP1 and IREB2/IRP2, respectively. Supernatants reacting specifically with ACO1 or IREB2 were validated by western blotting using extracts from wild-type versus ACO1- or IREB2-null mice.

### Protein extraction and Western Blots

Total liver protein lysates (*Aco1* and *Ireb2* strains) were prepared as follows: Livers were homogenised in 3 volumes (w/v) of RIPA buffer (150 mM NaCl, 50 mM Tris pH 8.0, 1% NP40, 0.5% Na-deoxycholate, 0.1% SDS and Roche complete protease inhibitors) using a Teflon homogenizer, incubated for 30 min on ice and centrifuged for 10 min at 16000 x g and 4°C. Supernatants were stored at −80°C. Protein concentrations were quantified using a BCA protein assay kit (Pierce). For the around-the-clock wild-type and *Smg6* mutant livers, total lysates from our previous study were used that had been prepared by the NUN (NaCl, Urea, Nonidet P-40) procedure, see [27].

Immunoblotting to PVDF membrane was carried out using iBlot 2 dry blotting system (Invitrogen). For the blots in Fig 4G and J, bi-directional diffusion transfer that results in two mirror image membranes per gel was carried out (protocol Ulrich Laemmli, personal communication). Briefly, PVDF membranes were placed on both sides of the gel and further sandwiched by 2 pre-wet Whatman filter papers and fibre pads. The stack was placed inside a transfer stack holder and submerged in a tank filled with transfer buffer (Tris-HCl 10 mM pH7.5, EDTA 2 mM, NaCl 50 mM and DTT 0.1 mM) and tightly sealed. The container was then placed in an incubator at 50°C for overnight diffusion transfer. Following transfer, membranes were blocked and incubated with primary and secondary antibodies in 5% milk/TBS-T. For IRP1 and IRP2 Western blots, undiluted crude hybridoma supernatants were used in primary antibody incubation. The dilutions for the remaining antibodies are as follows: anti-Transferrin Receptor (ThermoFisher 13-6800, 1:1000), anti-FBXL5 (ThermoFisher PA5-113529, 1:1000), anti-VINCULIN (Abcam 129002, 1:10000), rat monoclonal anti-ACO1 and anti-IREB2 (undiluted hybridoma supernatant from own preparation, see above); secondary antibodies: anti-rabbit HRP or anti-mouse HRP (Promega, 1:10000).

### RNA extraction and library preparation

#### Library preparation from footprints and total RNA

The libraries for ribosome profiling were generated as [6] with few modifications. Briefly, frozen liver tissues from individual mice were homogenized with 5-6 strokes using a Teflon homogenizer in 3 volumes of polysome buffer (150 mM NaCl, 20 mM Tris-HCl pH7.4, 5 mM MgCl_2_, 5 mM DTT, complete EDTA-free protease inhibitors (Roche) and 40 U/ml RNasin plus (Promega)) supplemented with 1% Triton X-100 and 0.5% Na deoxycholate. Liver lysates were incubated on ice for 10 min and cleared by centrifugation at 10000 g, 4°C for 10 min. Supernatants were flash-frozen and stored in liquid nitrogen. For each sample, 15 OD_260_ of liver lysate was digested with RNase I (Ambion) for 45 min at RT under gentle agitation. Digested samples were passed through pre-washed (4x with 700 μl polysome buffer for 1 min at 450 x g) Sephacryl S-400 HR spin columns (GE healthcare, currently Cytiva), and centrifuged for 2 min at 450 x g at 4°C. The resulting flow-through was mixed with Qiazol and ribosome-protected fragments (RPFs) were purified using miRNeasy RNA extraction kit (Qiagen) according to the manufacturer’s instructions. Total RNA was extracted from undigested liver lysates using Qiazol and miRNeasy kit in parallel to generate total RNA libraries. 1 μg of RPF and total RNA samples were depleted of ribosomal RNA using RiboCop rRNA depletion kit V1.3 (Lexogen), followed by purification with RNA Clean & Concentrator-5 (Zymo research). All the subsequent steps were performed according to Illumina’s TruSeq Ribo-Profile protocol with minor modifications as described in [69]. Size-selection of fragments at different steps was as follows: ribosome-protected fragments (RPFs) were size-separated on 15% urea-polyacrylamide gels and excised at a size range of 24-36 nt (monosome footprint). The cDNA fragments generated from RPFs and total RNA were separated on 10% urea-polyacrylamide gels and excised at a size range of 70-80 nt and 70-100 nt, respectively. The PCR-amplified libraries were size-selected on 8% native polyacrylamide gels. Monosome libraries were at ∼150 bp and total RNA libraries were at ∼150 to 170 nt. All libraries were sequenced on Illumina HiSeq 2500.

#### Ribosome profiling data analysis

Sequencing reads were mapped following overall the strategy described in [6]. Adapter sequences were removed from the sequenced reads using cutadapt (v3.5) with the following options: --match-read-wildcards --overlap 8 --discard-untrimmed --minimum-length 30 and reads were quality filtered using fastx_toolkit (-Q33 -q 30 -p 90). Trimmed read sequences were filtered by their size using an in-house Python script with following inclusive ranges: [26,35] for footprints, [21,60] for total RNA. Reads were then mapped to rRNA and tRNA databases to remove contaminants using bowtie2 (-p 2 -L 15 -k 20 --no-unal -q). Only reads that did not map to the above databases were mapped to mouse genome v38.100 using STAR (version 2.70f). Read counts of total RNA and RPF were normalised with upper quantile method of R package edgeR v3.4.2 [70]. Translational efficiencies (TE) were calculated on normalised reads as the ratio of RPF / mRNA for each gene per sample. TEs were log_2_-transformed and plotted for comparing means of replicates per time point for each genotype. K-Means cluster analysis was performed on the different conditions (averaged by replicates). Based on the Within-Cluster-Sum of Squared Errors, the number of clusters was fixed to 15 sets. Hierarchical clustering was then performed within individual clusters to assess proximity between genes of a given cluster.

### Liver explant preparation and bioluminescence recording

For explant preparation, male IRP1 or IRP2 KO and matched control animals from the same breedings (i.e., same genetic background) were euthanised following deep anaesthesia by isoflurane inhalation. Liver tissue was excised and placed immediately in ice-cold Hank’s buffer (Thermo Fisher Scientific). The outermost edges of the tissue were carefully excised in a sterile cabinet, and immediately placed on a 0.4 micron Millicell cell culture inserts (PICMORG50) in a 35 mm dish with phenol-free DMEM (Thermo Fisher Scientific, 11880028) containing 5% FBS, 2 mM glutamine, 100 U/ml penicillin, 100 μg/ml streptomycin and 0.1 mM luciferin. The parafilm-sealed plates were placed for recording in the LumiCycler (Actimetrics) at 37°C and 5% CO2.

### *In vivo* recording in the RT-Biolumicorder

Adult male mice, 12-20 weeks of age, carrying the genetically encoded circadian reporter allele *mPer2::Luc* [46] and deficient for *Aco1* or *Ireb2* were used for the RT-Biolumicorder experiments, along with controls, i.e. wild-type male mice from the same breedings/with the same genetic background and of the same age. The experimental procedure followed our previously published protocols [45]. Briefly, ALZET mini-osmotic pumps were filled with D-luciferin sodium salt and closed with blue-coloured flow moderators (ALZET) under sterile conditions. The pumps were activated at 37°C according to the manufacturer’s instructions, followed by subcutaneous, dorsal implantation. Before implantation, the dorsal area of the mouse at the site where the liver is positioned was shaved using an electric razor. The RT-Biolumicorder (Lesa-Technology) consists of a cylindrical cage equipped with a photomultiplier tube (PMT) allowing for recording of real-time longitudinal bioluminescence from single-housed mice. The data, which also contain locomotor activity traces, light and food access information, were saved as text files and analysed using the MATLAB-based “Osiris” software according to [45].

### Iron measurements from liver samples

Iron measurements were performed in principle according to the protocol of the Iron Assay Kit from Abcam (ab83366), with modifications serving to adapt to the specific properties of liver tissue (in particular the high lipoprotein content that is, moreover, rhythmic). 350 µl Iron Assay buffer and 20 µl 20% SDS were added to the liver pieces (50 mg) in 2 ml tubes for Precellys homogenisers (Ref. 432-0351, VWR/Bertin Instruments). Then, 3 small (1.4 mm; P000945-LYSK0A, VWR/Bertin Instruments) and 2 big (2.8 mm; P000927-LYSK0A, VWR/Bertin Instruments) ceramic beads were added to the tube. Tissues were homogenised using Precellys 24 (program: 5000 rpm, 2×20 sec, 9 sec pause between the two 20 sec runs). Lysates were cleared through centrifugation at 16,000 rpm, RT, in a microcentrifuge for 10 min. Supernatants were used in Iron Assay Kit according to the manufacturer’s protocol (Abcam, ab83366). 5 µl of 20% SDS were added to each well before addition of Iron Probe to overcome lipoprotein-caused turbidity. This led to the clearance of the samples, but also to precipitate likely from SDS and potassium. For this reason, all samples (including standard curve) were briefly re-spun and 100 µl of the supernatants were transferred to a fresh 96-well. The absorbance was quantified using a colorimetric microplate reader (at optical density 593 nm), and quantifications of iron states (total, ferrous, ferric) and further calculations were carried out as described in the supplier’s protocol.

### Statistical analyses

Statistical analyses were performed using GraphPad Prism. Unpaired t-tests and one-way ANOVA tests were applied for comparisons of two groups and multiple groups, respectively. Šidak’s post hoc correction was performed for multiple comparisons. All tests were two-tailed and represent a 95% confidence interval. P value of < 0.05 was defined as statistical significance.

## Declarations

### Ethics approval and consent to participate

All animal experiments were performed according to the cantonal guidelines of the Canton of Vaud, Switzerland, license VD3402.

### Consent for publication

Not applicable

### Availability of data and materials

All data needed to evaluate the conclusions in the paper are present in the paper and/or the Supplementary Files. Sequencing data have been deposited at GEO (GSE243134). The graphs using external data shown in Fig. 6 are based on the values submitted in the supplementary materials of the cited publications, i.e. Greco et al. [56] Data S1, https://www.science.org/doi/suppl/10.1126/sciadv.abi7828/suppl_file/sciadv.abi7828_data_s1_to_s9.zip, Atger et al. [7] Dataset S1, https://www.pnas.org/doi/suppl/10.1073/pnas.1515308112/suppl_file/pnas.1515308112.sd01.xlsx, Greenwell et al. [57] Table S2, https://ars.els-cdn.com/content/image/1-s2.0-S2211124719303948-mmc2.xlsx, Manella et al. [58] Supplementary Data 2, https://static-content.springer.com/esm/art%3A10.1038%2Fs42255-021-00395-7/MediaObjects/42255_2021_395_MOESM3_ESM.xlsx.

### Competing interests

The authors declare that they have no competing interests

## Funding

Swiss National Science Foundation (SNSF), NCCR RNA & Disease grant 141735 (DG)

Swiss National Science Foundation (SNSF) individual grant 179190 (DG)

## Authors’ contributions

DG, ABA, BG, HPN, and GK designed the research. HPN, GK, ESA, AL and GP performed the experiments. HPN, GK, ESA, GP, BG and DG analysed and interpreted the experimental data. LC and APA carried out bioinformatics analyses. GP, CT, IH and BG produced the new antibodies. DG, BG, HPN and GK wrote the manuscript. All authors read and approved the final version of the manuscript.

## Acknowledgements

We thank Matthias W. Hentze for knockout mouse strains; the Lausanne Genomic Technologies Facility for library preparation and sequencing; and Paul Franken and members of the Gatfield lab for comments on the manuscript. We thank Tomasz Martini, EPF Lausanne, for preparing Supplementary Fig. S2.

## Notes

### Competing Interest Statement

The authors have declared no competing interest.

### Summary of Updates

-Main new data: around-the-clock iron measurements. -Otherwise: Overall shortening and rewrite.

