## Supplementary Figures for "Diurnal control of iron responsive element containing mRNAs through iron regulatory proteins IRP1 and IRP2 is mediated by feeding rhythms"

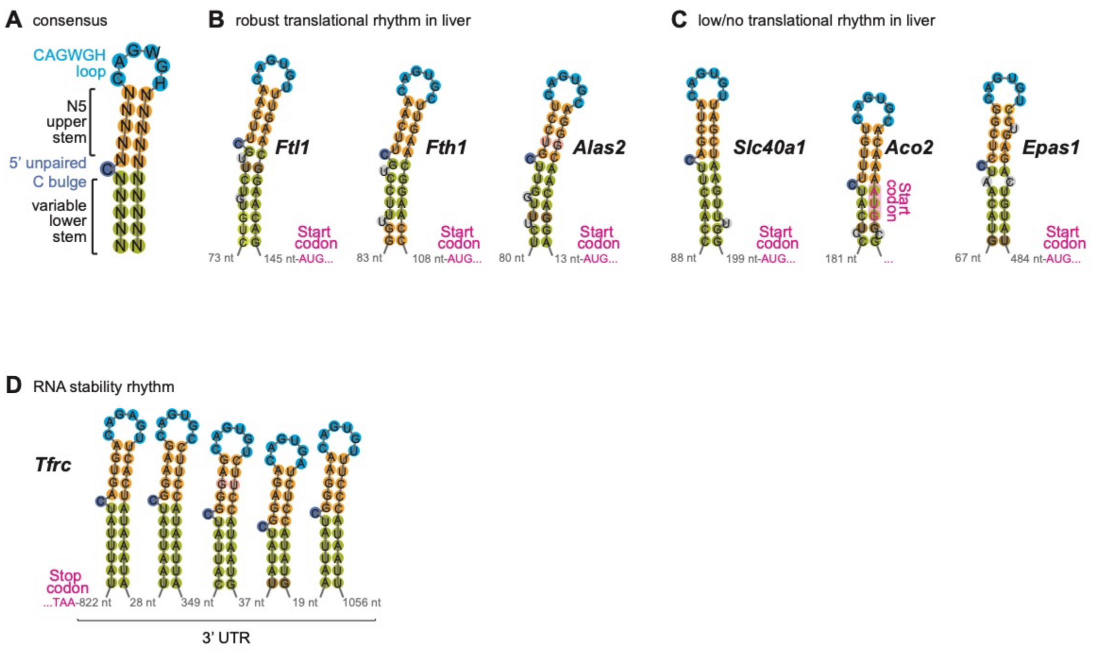


**Figure S1. Transcript- and tissue-specific rhythmicity of IRE-containing mRNAs.**

(**A**) Schematic of the IRE hairpin consensus sequence and structure. IREs are about 35 nt in length land are characterized by an apical 6-nt loop motif 5'-CAGWGH-3' (W = A or T and H = A, C or T), and two stem regions (N5 upper stem and variable lower stem) that are separated by a unpaired C bulge. Colour code depicts sequence conservation across IREs from high (blue) to medium (orange) and low (olive).

(**B**) Sequence and predicted structure, as well as relative position within 5' UTR, of the IREs of the murine transcripts *Ftl1, Fth1* and *Alas2* that are shown in main Fig. 1C. Color code as in (A).

(**C**) Sequence and predicted structure, as well as relative position within 5' UTR, of the IREs of the murine transcripts *Slc40a1, Aco2* and *Epas1* that are shown in main Fig. 1D. Color code as in (A).

(**D**) Sequence and predicted hairpin structures of the five IREs annotated within the mouse *Tfrc* 3' UTR. Nucleotide distances from the TAA stop codon, between individual haipins, and to the annotated transcript end are noted in grey. Color code as in (A).

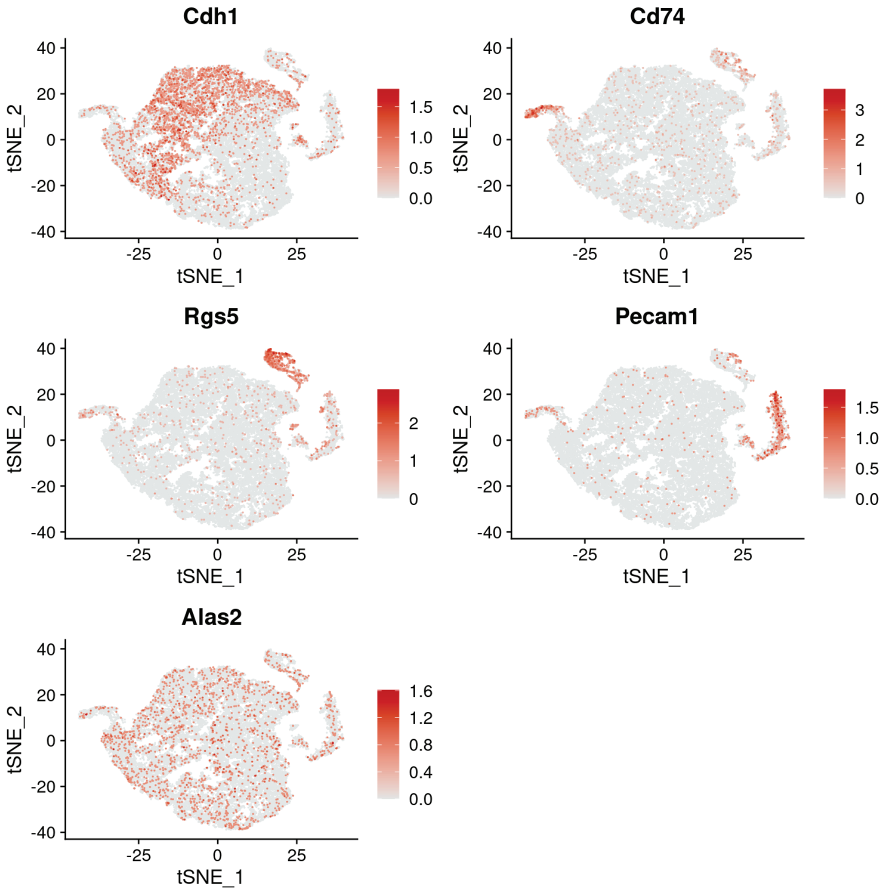


**Figure S2. Single-cell sequencing data validates hepatocyte expression of *Alas2*.** t-SNE plot of liver single cell RNA-seq analysis using data from Martini et al., bioRxiv 2023.10.07.561324. The large central group of cells are hepatocytes. As marker transcripts in the first 4 plots, *Cdh1* is specific for periportal hepatocytes; *Cd74* shows immune cells; *Rgs5* identifies stellate cells; *Pecam1* are endothelial cells. *Alas2* is lowly expressed everywhere, including across periportal (*Cdh1*-positive) and pericentral (*Cdh1*-negative) hepatocytes. Figure preparation acknowledgement: Tomasz Martini, EPF Lausanne, Switzerland.
